# Separate domains of the *Arabidopsis* ENHANCER OF PINOID drive its own polarization and recruit PIN1 to the plasma membrane

**DOI:** 10.1101/2024.03.11.584374

**Authors:** Michaela S. Matthes, Nicole Yun, Miriam Luichtl, Ulrich Büschges, Birgit S. Fiesselmann, Benjamin Strickland, Marietta S. Lehnardt, Ramon A. Torres Ruiz

## Abstract

The *Arabidopsis* ENHANCER OF PINOID (ENP) protein and the AGC-kinase PINOID (PID) synergistically impact on polarization of the auxin transporter PIN-FORMED1 (PIN1) required for plant leaf and flower organ development. ENP offers a PID-independent input for PIN-polarity since *enp pid* double mutants lead to cotyledon- and flower-less plants in contrast to *pid* single mutants, which develop cotyledons and abnormal albeit fertile flowers. This indicated that ENP, which depicts a similar polar localization as PIN1, is a potential interactor of PINs especially PIN1.

Here we show that the modular structure of ENP predicted by AlphaFold separates the capability for its own cellular polarization and its function linked to polar PIN1 activity. The anterior part of ENP is subdivided into three structured domains. They are supportive and/or essential for cellular polarity. In contrast, the C-terminus, which is an intrinsically disordered region (IDR), is completely dispensable for polarity but essential for ENP-mediated PIN-function. FLIM-FRET shows ENP to be closely associated with the plasma membrane and its IDR to significantly interact with PINs. Moreover, the modification status of two prominent phosphorylation sites in the IDR determines ENPs stability and its capability in supporting PIN1. Our results show ENP to be an element in the assumed PIN-multiprotein complex and explain its impact on PID-independent PIN1 activity.

## Introduction

The plant hormone auxin works as a concentration-dependent signal molecule controlling various plant developmental processes [1, 2]. During embryogenesis local auxin concentrations (auxin maxima) are read out to induce the generation of root vs. cotyledons, the embryonic leaves [3]. They also impact on cotyledon number and shape [4]. During adult development, auxin controls processes such as the generation of leaf and flower primordia [3]. Auxin maxima are organized by a system of auxin influx and efflux carriers. In the cytosol, the main auxin indol-3-acetic acid (IAA) is a charged, membrane-impermeable molecule making its transport predominantly dependent on efflux carrier proteins [5]. The most important efflux carriers are the closely related plasma-membrane (PM) integral PIN-FORMED proteins (PINs) [3, 6]. PINs have been shown to be organized as homodimers, which export auxin via a transport mechanism described as elevator-like, their localization indicating the transport direction [7–9]. Correspondingly, PINs are apically polarized in epidermal cells of (aerial) organ primordia, while they adopt a basal orientation pointing towards the root tip in inner tissues [3]. The basal orientation of PIN1 is controlled by GNOM [10]. The apical polarity of PINs is affected by numerous factors. Among these, the site-specific phosphorylation of PINs by different kinases counteracted by phosphatases [11] is essential although it is debated whether phosphorylation *per se* or a complex temporal/spatial pattern of dynamic de-/phosphorylation determines PIN polarity [1, 2, 12]. The AGCVIII family kinase PINOID (PID) works as a developmental switch for PIN1 polarity and has a crucial role in shoot development [13]. *Pid* single mutants generate pin-like inflorescences, like *pin* single mutants, but also stems with abnormal but fertile flowers providing seedlings with two or three cotyledons [14, 15]. This has been attributed to the observation, that *pid* mutants retain some apically polarized PIN1 in the epidermis of cotyledon primordia [16]. In contrast, double mutants of *PID* and *ENHANCER OF PINOID* (*ENP*) completely lack flowers as well as cotyledons, which correlates with a shift of PIN1 to lateral and basal epidermal cell poles [16]. A number of mutants in *pid* background leading to (cotyledon) abnormalities uncovered additional genes involved in these processes. This concerns *pin1* itself and genes required in auxin biosynthesis, the Hippo signalling pathway and endosomal sorting [17–21].Together, they suggest the presence of another rather PID-independent input, which contributes to organogenesis.

ENP, also named MACCHI-BOU4/MAB4 [22] or NAKED PINs in YUCCA1/NPY1 [23] and four additional proteins called MELs (<u>M</u>AB4/<u>E</u>NP/NPY-<u>L</u>ike) display similarity in the N-terminal and central domain with the NON-PHOTOTROPIC HYPOCOTYL3 (NPH3) protein while their C-termini exhibit considerable divergence [18, 22–24]. The expression pattern of *ENP* (*MAB4; NPY1*) vs. *MELs* is complementary. *ENP* is prominent in its epidermal expression while *MELs* are mainly expressed in internal tissues [18, 22]. ENP’s apical and MEĹs mainly basal [24] cellular polarities overlap with the known polarities of PINs. This and the phenotypes of *enp pid* and multiple combined *ENP/MEL* mutants suggested a role in auxin transport including (genetic and/or physical) interaction with PINs. In fact, recently in vitro pull-down essays of MEL1 with PIN2 and ENP(MAB4) with PIN2 indicated physical interaction. In addition, the interaction of MEL1 with PIN2 was shown by FLIM-FRET [25].

In this study, we have focussed on the molecular characterization of ENP and its contribution to PIN1 activity. Here we show, that ENP’s architecture consists of separated modules required for two different functions: first, the capability of ENP for its own polar localization in the cell; second, the support of PIN1 leading to restoration of the *pid* single mutant phenotype in *enp pid* background (“*enp pid* rescue”). For convenience the former function is termed (ENP) “polarity” and the latter (PIN-supporting) “functionality” in the following text. The N-terminal and in particular the central region covers ENP’s capability for apical localization. Although the integrity of these parts is necessary for the overall function of ENP, they alone cannot support PIN1. In contrast ENP’s C-terminus, an intrinsically disordered region (IDR), is dispensable for ENP localization but absolutely essential for its function recruiting PIN1 to apical plasma membrane domains in the epidermis. A comparison with MEL4/NPY4, which largely lacks a C-terminal part, shows that its similarity to ENP is sufficient for the same polar capabilities (polarity) but not to replace ENP in its PIN1 supporting function (functionality). Furthermore, our data show that the functional strength of the C-terminus increases depending on the integrity and modification of at least two known phosphorylation targets. FRET analyses using PIN2, the structural and functional homolog of PIN1 in the root epidermis, shows that ENP interacts with the cytosolic loop of PINs. The same technique shows that ENP is closely associated with the PM.

## Results

ENP is a protein of 571 amino acids (aas) with a modular architecture (Fig. 1; SFig.1). The N-terminus (from aa1 to aa132) contains a BTB/POZ domain (aa29-aa132), which is a conserved protein-protein interaction motif originally found in poxviruses, mice and *Drosophila melanogaster* involved in a variety of functions [26 and references therein]. X-ray crystallography data have identified tertiary/structural similarity while there is little sequence similarity between different protein families [26]. ENP is a member of plant-specific BTB-NPH3 proteins, whose N-terminus is predicted by AlphaFold to have numerous α-helices and β-sheets with high likelihood as quantified by a residue confidence score called predicted Local Distance Difference Test value (pLDDT) [27, 28] (Fig. 1 H, J; SFig.1). These are per residue confidence scores scaled between 0 and 100 indicating how well the predicted structure would agree with the experimental structure. In many BTB-proteins the N-terminal BTB/POZ-domain is followed by a linker region, which connects to the following domain [26]. In ENP, the region from aa133 to aa210 is tentatively designated as “linker” and the larger region reaching to aa470 as “central core” with portions of alternating high and low similarity to NPH3 (NPH3_1 to NPH3_3) [26]. The “central core” contains α-helices of variable length interrupted by only one region of unpredictable structure (aa185 to aa205). The adjacent C-terminal region (aa 471 to aa571) is quite diverse between ENP and all MELs; in MEL4 it is almost completely missing (Fig. 1 G-J; SFig.1). By phylogenetic analysis the anterior parts of ENP, MELs and NPH3 show high similarity while the C-termini are highly dissimilar. This division also exactly overlaps with the structural prediction by AlphaFold [27, 28], which indicates intrinsic disorder from aa471 to aa571 by low pLDDTs scores, which have been shown to be suitable predictors for intrinsic disorder of protein regions [29, 30].

**Fig. 1:**
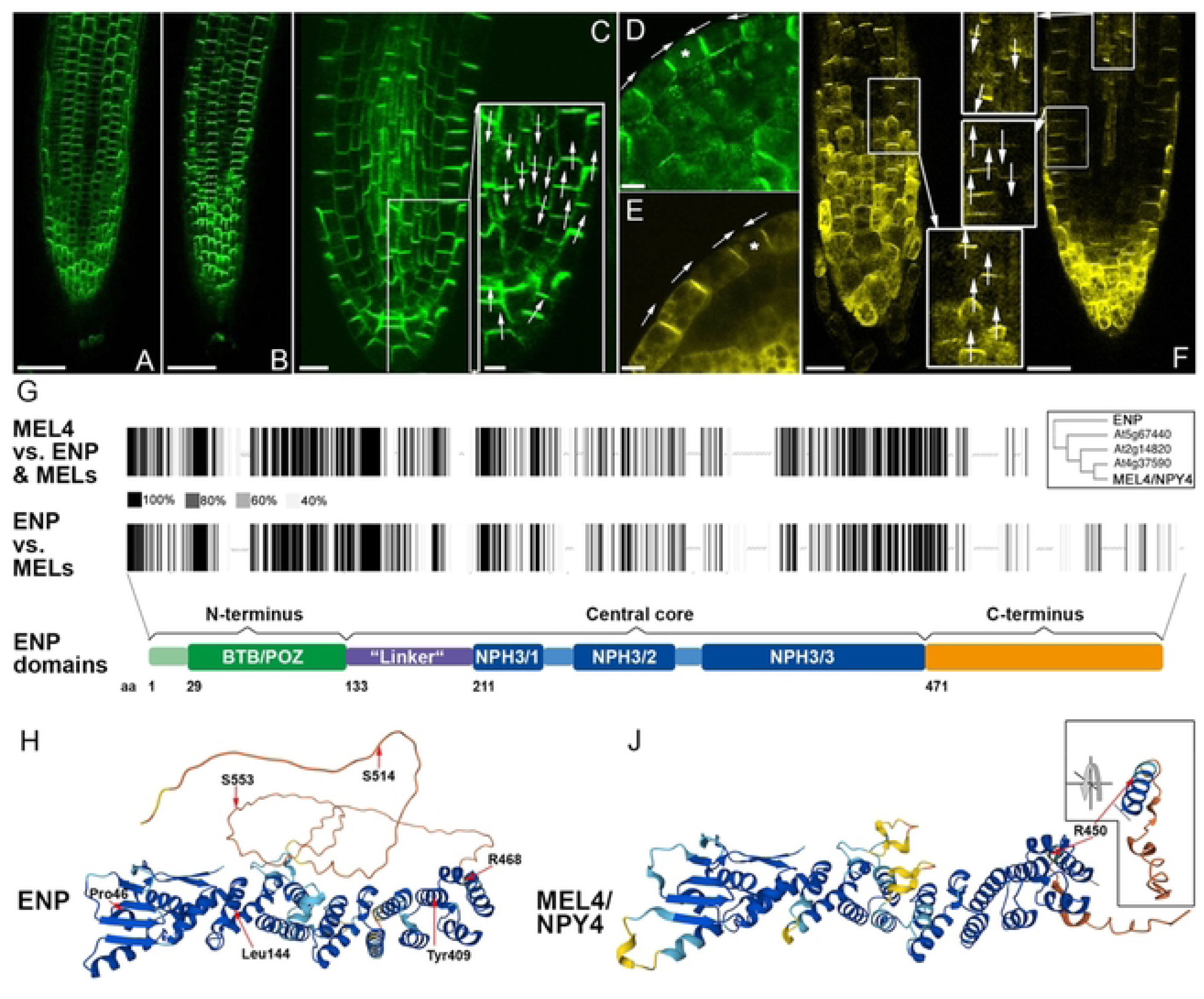
Cellular polarity and structure of ENP and MEL4. A) EGFP-ENP (focus on epidermis, *35Sp:EGFP-ENP).* B) ENP-GFP6 (focus on epidermis, *35Sp:ENP-GFP6).* C) ENP-GFP6 (focus on internal tissues). D)EGFP-ENP in embryo cotyledon. E) MEL4-EYFP in embryo cotyledon. F) MEL4-EYFP in the seedling root. Left: focus on the epidermis, framed region magnified in the bottom inset; right: focus on the internal tissues/stele, framed regions magnified in the top and middle insets. G) Protein similarity schemes. The upper bar code compares MEL4 with ENP and the proteins MEL1-3; the lower ENP with MEL1-4; given are gray to black lines with degree of similarity indicated. Note the C-terminal region of ENP vs. the much shorter terminal region of MEL4. The inset depicts a similarity tree between these proteins based on ClustalW. H) AlphaFold-predicted structure for ENP (Jumper et al., 2021; Varadi et al., 2021). J) AlphaFold-predicted structure for MEL4/NPY4. Inset shows turn of the terminal region to highlight position of amino acid R450 (the analogue of R468 in ENP). Studied amino acid residues are indicated. See text Color code for the per residue confidence metrics in H, J) dark blue: pLDDT>90, light blue: 90>pLDDT>70, yellow: 70>pLDDT>50, orange pl DDT<50. Scale bars: 50µM in A, B; 10µM in C; 5µM inset in C, D, E; 20µM in F. White arrows indicate polar orientation of EGFP-ENP, ENP-GFP6 and MEL4-EYFP respectively. Star in D and E shows the terminal cell. Here polar ENP and MEL4 respectively, face each other in opposite cells.

### ENP displays tissue dependent apical and ectopic basal polarity

ENP and MELs have significant similarity in the N-terminus and central core (Fig.1; SFig.1). While ENP is expressed and apically localized in epidermal cells, *MELs* are mainly expressed in cortex and stele cells where they are basally localized. We were interested to analyse whether ENP is capable to polarize in other than epidermal cells. To this aim, “wild-type” i. e. full-length ENP cDNA-GFP constructs driven by the *35S* promoter were analysed in seedling roots and embryos (*35Sp:EGFP-ENP, 35Sp:ENP-mGFP6;* Fig. 1A-D; SFig.2). Our interest focussed mainly on seedling roots and embryos. The orientation of ENP was apical in epidermal tissue, while it was basal in internal tissues (Fig. 1A-D). Expression of ENP itself seemed not to be affected in cells with altered/interrupted auxin transport as given in *enp pid* double mutant embryos. *In situ* hybridization showed epidermal *ENP* mRNA signal as in wild-type (SFig.3) [22]. We conclude, that ENP possesses information for polar localization in all cells. The readout of this information differs in epidermal vs. internal tissues leading to apical vs. basal localization, which essentially overlaps with that of PIN1 and PIN2 in accordance with CoIP results for ENP/MAB4 and PIN2 [25].

**Fig. 2:**
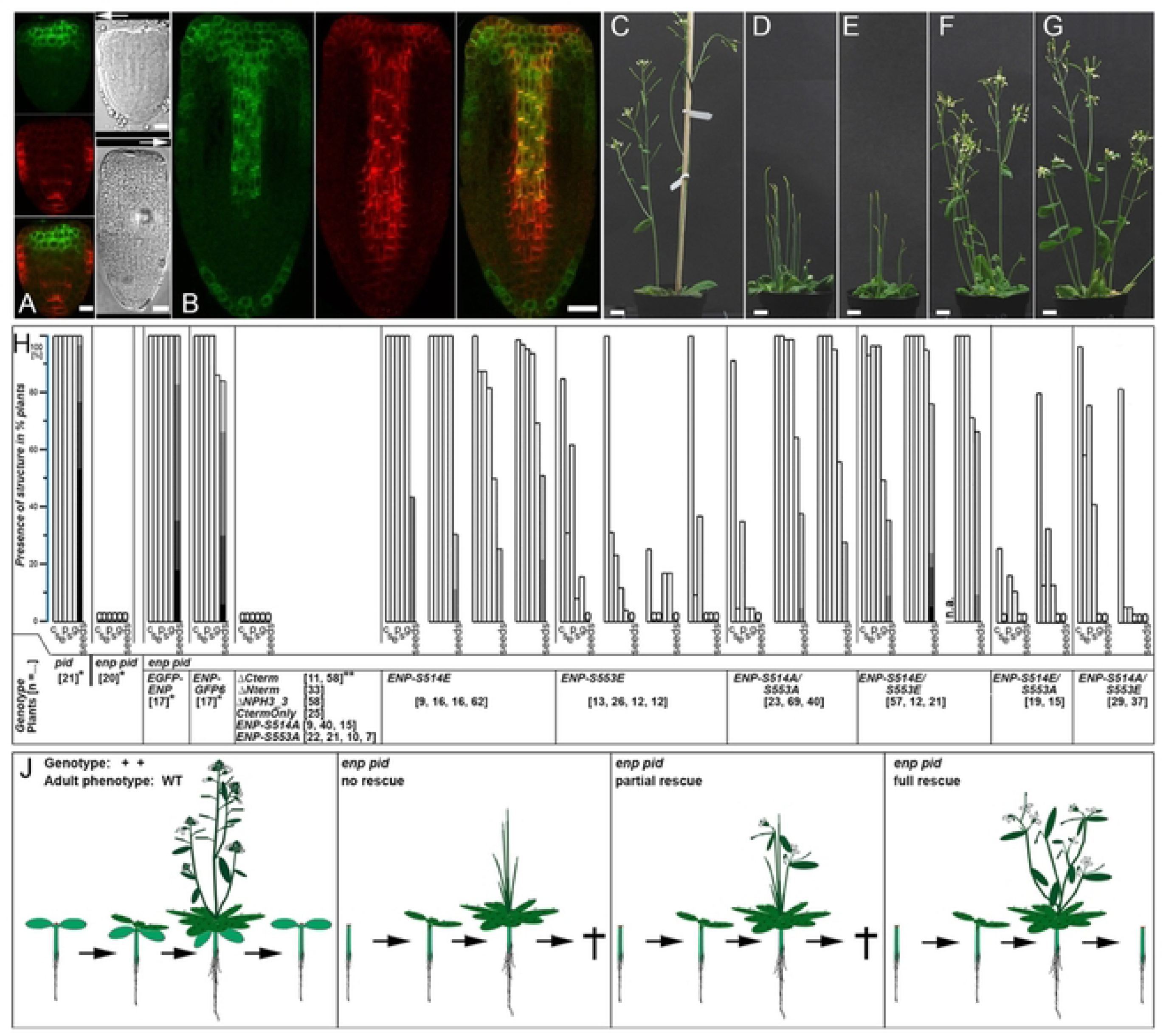
Timing of ENP expression and the rescue of *enp pid* double mutants. A) Late 35Sp-driven ENP-GFP signal in the central (not lateral) apex region of *enp pid* heart stage embryos. B) 35Sp-driven ENP-GFP signal proceeds to but does not reach the root tip of *enp pid* torpedo stage embryos. The extension of PIN1 is shown for comparison (PIN1 AB-Cy3-staining; left: GFP; middle: Cy3; right: merger; see SText). C-G) always *A. thaliana Ler* ecotype background with C) wild-type *plant,* D) *enp pid* homozygous plant, E) *enp pid* homozygous *plant* with non-rescuing construct, F) *pid* homozygous *A. thaliana Ler* ecotype, G) *enp pid* homozygous plant with rescuing construct. Scale bars in A: 10µM, B including bottom left: 20µM, C-G: 1 cm. H) Frequencies organs generated and pedigree produced by independently transformed lines in homozygous *enp pid* plants (constructs indicated; see text and SFig. 2). The data for few representatives of *enp pid* and *pid* homozygous mutants without constructs are given (many more have been inspected along the course of this study with the same outcomes). Number of assessed plants per independent transformed line are given in brackets. Abbreviations C: cauline leaves or bracts, Se: sepals, P: petals, S: stamina, G. gynoecia, and seeds. Seed production is scored as plants with 1-25 (light gray), 26-100 (medium gray), 101-200 (darker gray) and> 200 (dark gray) seeds. J) Outer left: one full wild-type life cycle; inner left: *enp* pid-plants and *enp pid­* plants with non-functional constructs do not produce organs except rosette leaves; inner right: phenotype of plants with constructs competent for partial rescue (= generate bracts/cauline leaves and flower organs but no seeds; outer right: constructs competent for full rescue(= bracts/cauline leaves, flowers and seed production) thus producing cotyledon-less *enp pid* seedlings again. All plants with constructs cannot generate cotyledons due to late *35Sp* activity in the embryo!

**Fig. 3:**
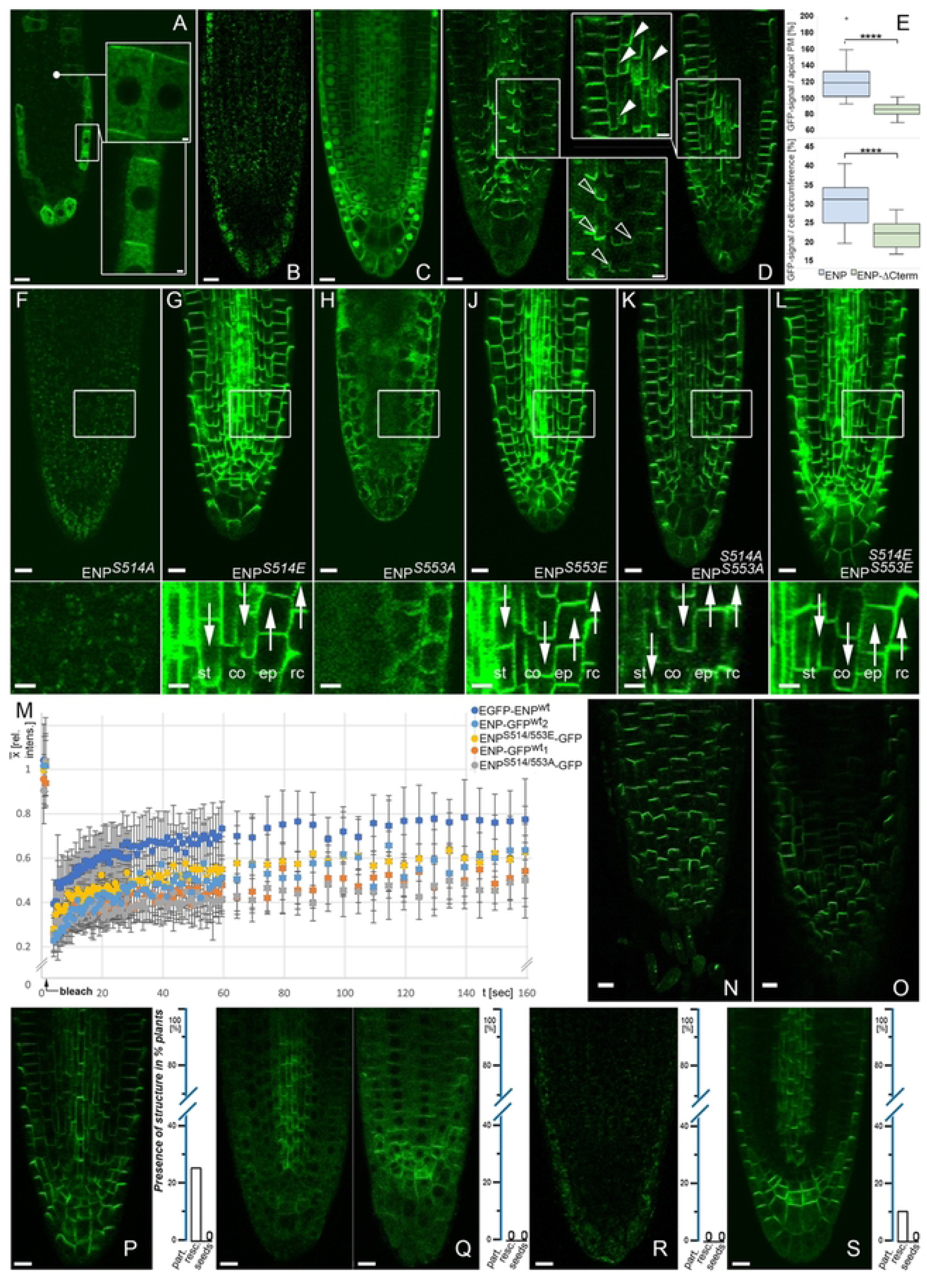
Properties of deletion and point mutation ENP constructs. A-E) Deletion constructs. A) *35Sp:ENPt:.Nterm-GFP6 construct.* Insets: magnifications. Note, the inset at the top is a magnification of the same specimen in the indicated region at slightly different focus. B) *35Sp:ENPt:.NPH3_3-GFP6* construct. C) *35Sp:ENPCtermOnly-GFP6* construct. D) *35Sp:ENPt:.Cterm-GFP6* (left) in comparison with full length *35Sp:ENP-GFP6* (right). Insets: open vs. filled arrowheads point to the different lateral extensions (“smile”) of ENP-t:.Cterm vs. ENP full length at the PM respectively (cortical and stele cells compared). E: I-Test for (lateral) extension of ENP-t:.Cterm vs. ENP at the PM (p<0,0001; see SText). F-L) ENP-GFP localisation of point mutations in critical C-terminal sites S514 and S553. Point mutations in the constructs indicated. White framed regions magnified at the bottom of each image. Auxin flux (as derived by polarity of ENP-GFP) indicated by white arrows in stele (st), cortex (co), epidermis (ep) and root cap (re) cells. All constructs with *35Sp* and *GFP6.* Brightness and contrast in F and H elevated to visualize residual GFP signal. M) Mobility of ENP S514/S553 double mutants compared to wild-type ENP as analysed by FRAP. Comparison of wild-type and mutant ENP transformants (indicated). Data generated from FRAP-analysis of at least three independent transformants each. N, 0) Polar localisation of ENPS514E/S553A (N) and ENPS514A/S553E (0). P-S) Selected plants harboring constructs with point mutations in the N-terminal, linker and central core region. P) *35Sp:ENP“’^46^T-GFP6* construct. Q) *35Sp:ENF>l^1440^-GFP6* construct, stele in focus; (left), epidermis in focus (right). R) *35Sp:ENPY^4^o^9^E_GFP6* construct. S) *35Sp:ENPv^4^o^9^A_GFP6* construct. Plants with *35Sp:ENP”’^467^-GFP6* (P) and *35Sp:ENPY^409^A-GFP6* construct (S) were capable of partial rescue. The former could generate bracts/cauline leaves and flower structures; the latter only bracts/cauline leaves. Absence of rescue capability and percentages of partial rescue respectively are indicated in the scale to the right. Scale bars A-D, F-L, N, 0 and P-S: 1OµM, insets in A: 1 µM, D, F-L: 5µM.

### The related MEL4/NPY4 shows the same cellular polarities as ENP

Among the *MEL1-4* (*NPY2-5*) genes, we selected *MEL4/NPY4* (hereafter MEL4) for comparison with *ENP*. Both share some interesting features but also display important differences. Most important is their similarity in the N-terminal and middle regions and the fact that MEL4 almost completely lacks a C-terminus in comparison to ENP and the other MELs/NPYs (Fig. 1G-J; SFig.1). Furthermore, while *ENP* is expressed in the epidermis, *MEL4* is expressed in the stele where the protein displays basal polarity [18, 24] (Fig. 1A-F). Again, both proteins are internalized to the cytosol upon phenylboronic acid treatment and thus share a similar response to this chemical affecting PM association (SFig.4) [31, 32].

**Fig. 4:**
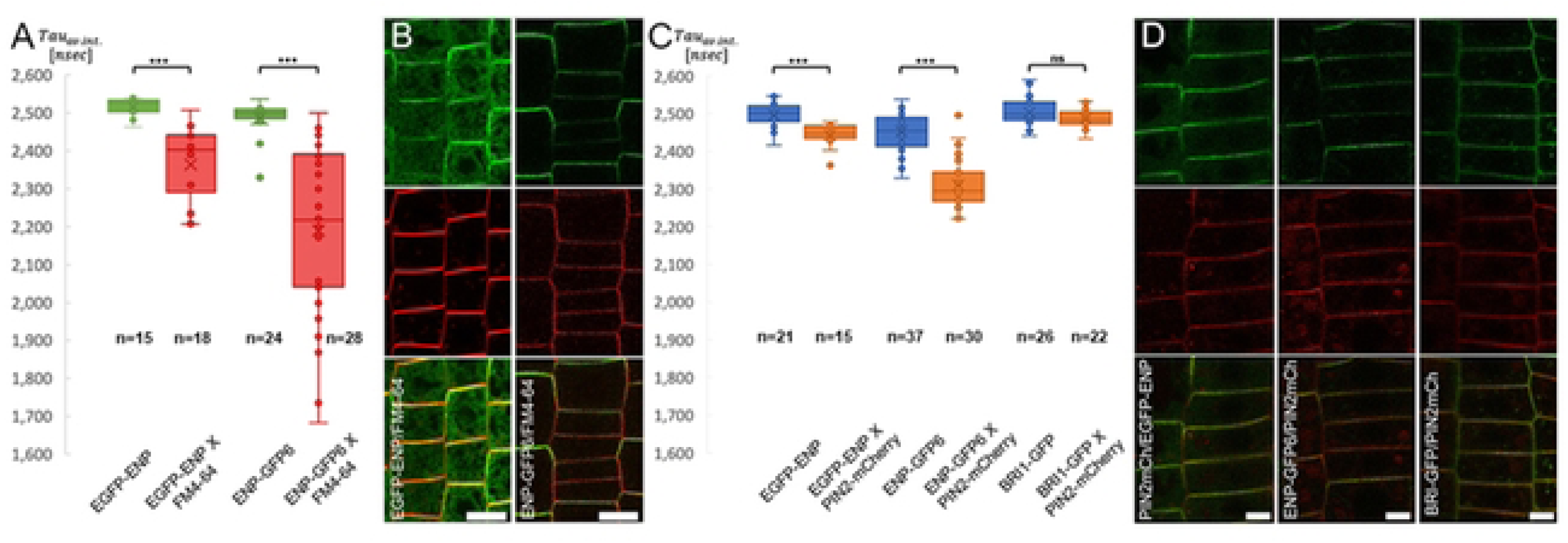
FLIM-FRET analyses of ENP-GFP with FM4-64 and PIN2-mCherry as acceptors. A, B) ENP constructs with/without FM4-64 (as indicated). A) ENP with N-terminally and C-terminally fused GFP display significant FRET in presence of FM4-64. B) Representative images showing GFP and FM4-64 fluorescence (separated lop and middle) and the merger (bottom). C, D) ENP and BRI constructs in absence/presence of PIN2-mCherry (as indicated). C) The TAUaverage intensity values of the donor only and donor with acceptor combinations as indicated are given. Differences are significant (p<0.0001) or not significant (p>0.05) with one-tailed t-Test (see SText). D) representative fluorescence images of EGFP-ENP X PIN2 (left), ENP-GFP6 X PIN2 (middle) and BRI-GFP X PIN2 (right) seedlings showing GFP fluorescence (lop), mCherry fluorescence (middle) and the merger (bottom). Scale bars in B: 10µM, in D: 5 µM.

We tested ectopic expression of *35S*p-driven *MEL4-EYFP* fusion in late wild-type embryos and seedling roots. *MEL4* constructs showed weaker fluorescence in comparison to *ENP* constructs but the protein clearly adopted a basal orientation in inner tissues and an apical localisation in the epidermis (Fig. 1E, F).

Thus, MEL4 also possesses sufficient information for apical and basal polarity in epidermal and inner tissues respectively. The information for apical vs. basal PM localization of ENP and MEL4 (and the other MELs/NPYs) is likely encoded in their N-terminal and/or central core.

### A genetically based system to efficiently assess polarity vs. functionality of ENP constructs in *enp pid* plants

For a molecular characterization of ENP, we wanted to investigate the impact of its domains on cellular polarity and functionality in terms of rescue of the flower-less phenotype of *enp pid*. Since the application of pyro-sequencing for the assessment of transgenic and mutant/wild-type genotypes proved to be unsuitable (SFig. 5), we established a genetically based bio-essay, which ensured a *pid enp* double mutant background and simultaneously allowed to assess the cellular localisation and the developmental functionality of constructs. Our approach implements, that ENP is required for both early (embryonic) and late (flower) developmental stages (Fig.2) [16]. Furthermore, constructs driven by the 35S promoter are known to be expressed only late in embryonic and then from early on in adult plant development. As a corollary, the onset of *35Sp*-driven *ENP* expression prevents the rescue of cotyledon development (SFig. 6). A comparison with PIN1, driven by its endogenous early promoter, illustrates this point in *enp pid* embryos. PIN1 is present in the whole embryo from early on while ENP is lagging behind in the apex (laterally where cotyledon primordia would initiate) and in the root (Fig.2 A, B). Thus, the analysis of an ENP construct comprised the following steps. First, selection of antibiotic resistant cotyledon-less seedlings assessed the *enp pid* homozygous background genotype and the presence of the corresponding ENP construct. Second, cellular localization of the GFP-fused protein could be assessed by CLSM in *enp pid* or (resistant) wild-type siblings. Third, resistant *enp pid* seedlings were grown to maturity. The absence or presence of any organs on stems (bracts, cauline leaves, flower structures) and pedigree was inspected and scored as no, partial or full functional activity of the introduced ENP version (Fig. 2H, J). Both mutants (*enp-1* and *pid-15*) used in this study are strong alleles in the *A. thaliana* L*er* ecotype background [16, 22].

**Fig. 5:**
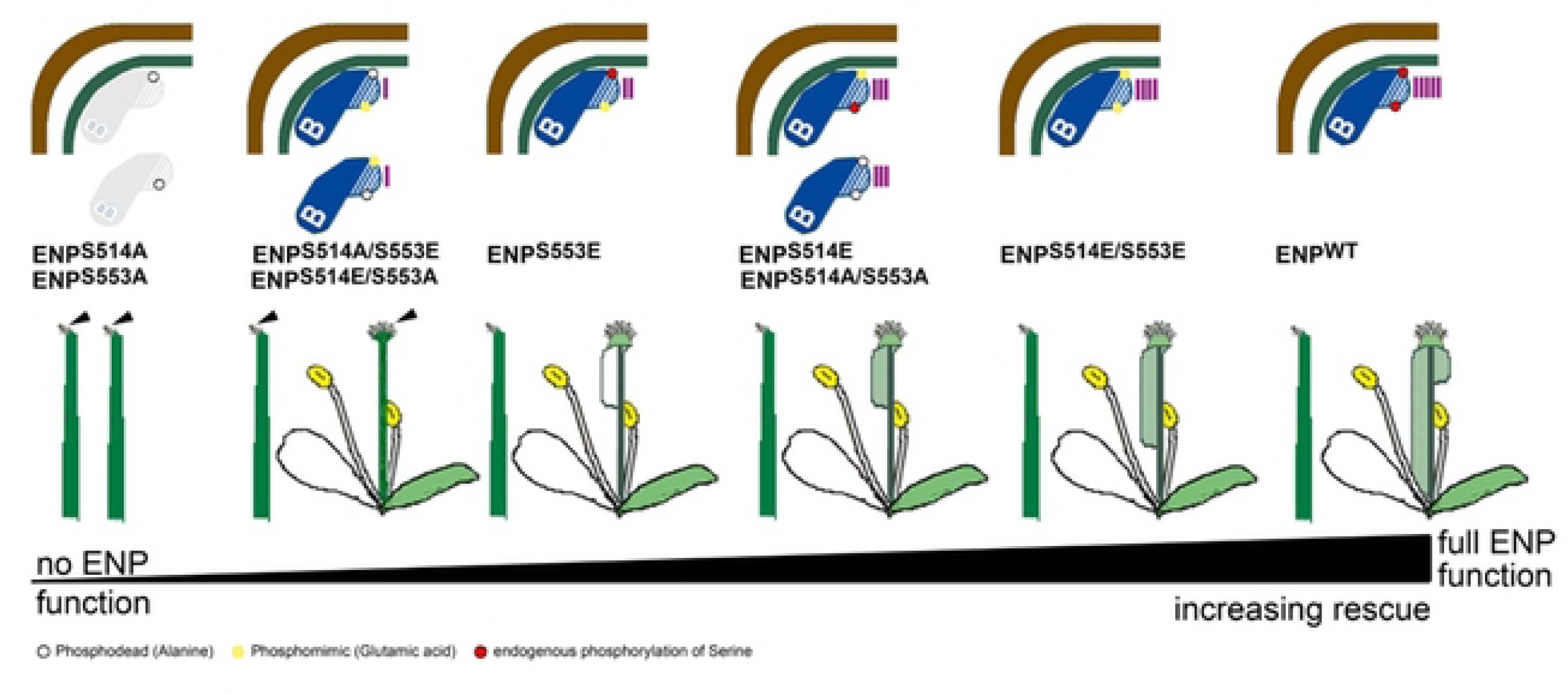
Strength of ENP C-term phosphomimic variants in *enp pid* rescue. Indicated is the progressive improve of flower development in *enp pid* plants by **ENP** constructs with modified C-termini, which progressively approach optimal interaction with PINs (vertical bars). Starting with unstable single phosphodead modifications at S514 or S553 (open circle), the rescue of *enp pid* improves gradually from left to right. Note, the double phosphodead construct is in the middle of this series. The grade of rescue is symbolized by flower-less, blind stems and irregular flowers without gynoecia (no carpels), gynoecia with empty carpels (white, no seed development) and gynoecia with increasing carpels tissue (which carried increasing seed set as given in Fig. 2H). All classes develop blind stems though with decreasing frequency towards the ENPwr constructs in the *enp pid* double mutant on the right. These latter plants are close if not phenotypically identical to *pid* single mutants. The progressive improve of flower development was paralleled by an increasing number of stems carrying bracts/cauline leaves (see Fig. 2H). Note that stigmatic papillae could develop even on blind stems and tissue replacing gynoecia (arrowheads). For details see text.

**Fig. 6:**
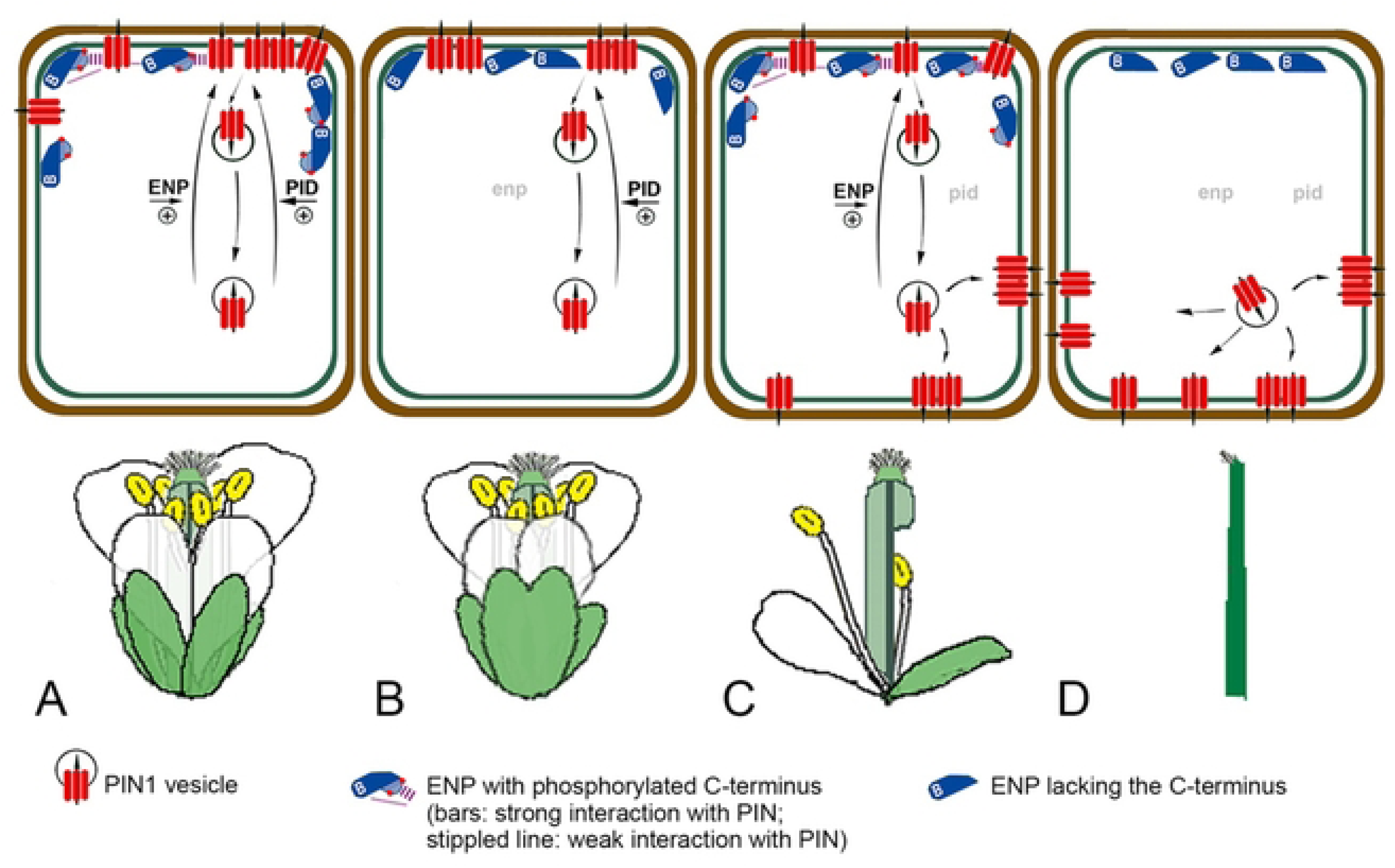
The independent impact of ENP on apical localization of PIN1. The scheme models the contribution of ENP and PIO activity on PIN1 polarity and development of flower structures (other factors are not included, see text). A) In wild-type both activities ensure optimal polarity of the PIN1 protein population. PIN1 vesicles are efficiently recruited to the apical end of epidermal cells. The interaction of ENPs C-terminus with PINs contributes to this polarity. B) This contribution is lost in the truncated *enp-1* single mutant although the truncated protein is still apically localized. PIO activity stillsustains polarization of sufficient but less PIN1 molecules for the development of most organs except sepals which are often fused. C) In *pid* single mutants the major impact on PIN1 polarity is absent but ENPs contribution sustains residual flower development including seed set. The correct phosphorylation of ENP (C-terminus) ensures optimal activity. PIN1 is increasingly localized to basal and lateral plasma membrane regions. 0) In *enp pid* double mutants both activities are absent and PlN1 is only laterally and basally localized (default basal GNOM transport). Leaf and nower organs on stems (as well as cotyledons) are completely absent in these plants.

### ENP but neither MEL4 nor MEL4/ENP-Cterminus domain swaps rescue *enp pid* double mutant *Arabidopsis thaliana*

Cotyledon-less seedlings carrying (E)GFP fused to either the N– or C-terminus of ENP were capable to fully rescue the *enp* mutation. These plants, developed rosette leaves, stems, leaf-like structures and *pid*-like flowers. Notably, they produced 100% cotyledon-less progeny, thus representing a new cotyledon-less plant population (Fig.2 C-J). The amount of progeny was often comparable to the *pid* single mutant.

We then tested whether MEL4 could functionally replace ENP. However, none of the *enp pid* plants carrying *35Sp:MEL4-EYFP* (n = 63) produced any leaf-like structures on stems. Since MEL4 almost lacks a C-terminus in comparison to the other MELs, we tested whether the addition of the ENP C-terminus could convert MEL4 to a (partially) functional version. We generated two constructs (see SupplText; SFig.2). One version fused the ENP C-term from aa471 to aa571 to the MEL4 protein fragment aa1 to aa452 (*35Sp:MEL4-ENPC term_long-GFP6*). The other was the addition of slightly shorter ENP C-terminal part (aa500 to aa571) to compensate for the fusion of the complete MEL4 protein (aa1 to aa481) (*35Sp:MEL4-ENPC term_short-GFP6*). Both variants did not lead to any rescue (n = 14 and n =34 respectively). Thus, MEL4 is not sufficiently similar to ENP in order to rescue the *enp* mutation even when extended with C-terminal parts of ENP.

### ENPs central core region is required for polar localisation

Next, we tested the significance of the N-terminal region up to the end of the central core for cellular polarity. To this end we first deleted the N-terminus from aa1 to aa53, which also deletes 25 aas of the BTB/POZ domain (ENP-ΔNterm construct; SFig.2). These plants display predominantly cytosolic distribution of the GFP signal. However, careful inspection revealed residual GFP signal with apical polarity in epidermal layers (Fig. 3A). This construct could not even partially rescue *pid enp* (Fig. 2).

We then introduced deletions starting from the C-terminus. The first deletion covered the region from the last aa571 to aa369 (ENP-ΔNPH3_3; SFig.2). This deletion resulted in irregular, distributed cytosolic signal, sometimes in patches, no localisation at the PM (Fig. 3B) and no rescue (Fig. 2).

We analysed two additional constructs. The ENP-CtermOnly construct represents the complete C-terminal intrinsic disordered region, which consist of 100 aas in length (SFigs. 1, 2). It produced an abnormal pattern with strong localisation in the nucleus and the PM in general and less localisation in the cytosol (Fig. 3C). No rescue could be observed (Fig. 2).

The ENP-ΔCterm construct spans aa1 to aa470, the domains for which AlphaFold predicts structure. This ENP-ΔCterm deletion mimicked the original *enp* allele (*enp-1*) [16, 22], which converts aa468 (R) into a STOP codon (Fig. 1H; SFig.1). R468 lies at the end of the last alpha-helix predicted by AlphaFold and the last region of similarity between ENP and all MELs (Fig. 1G-J; SFig1). A homologous residue is also found in MEL4 in position 450, with similar AlphaFold confidence metrics (Fig. 1J; SFig. 1). The next aas up to aa571 have only very low similarity to MELs. MEL4 almost lacks this part completely. Nevertheless, ENP-ΔCterm plants showed the same cellular polarity pattern as the full length ENP with considerable signal strength. However, detailed inspection showed that the distribution of this construct was somewhat restricted. Quantitative analysis of GFP-fluorescence of full length ENP vs. ENP-ΔCterm constructs showed that the former displayed stronger extension (“smile”) to lateral sites (Fig. 3D, E; SFig. 7). Notably, the plants analysed did not show any rescue (Fig. 2). We conclude that ENPs N-terminus and especially the central core contains sufficient information for polar localisation. Actually, ENP retains polarity even when a considerable part of the N-terminus is deleted.

### Phosphomimetics prove Ser^514^ and Ser^553^ in the C-terminal IDR to be critical for ENPs functional capability

The deletion constructs tested showed that the N-terminus and even more the central core of ENP is required for polarity while the C-terminus (aa471-aa571) *per se* is not. Conversely, the N-terminus and central core alone are not at all capable to rescue *pid enp*. For this the C-terminus is obviously essential.

Besides intrinsic disorder, ENPs C-terminus thus displayed another hallmark of IDRs: functionality [33]. Moreover, IDRs have been reported to frequently harbor (functional) phosphorylation sites, especially serines and threonines [34–36]. For ENP, the phospho-proteome database (PhosPhAt4.0 database; https://phosphat.uni-hohenheim.de/) lists serine target sites, the two most prominent localized in the C-terminus. These were especially conspicuous in three aspects: data quality, abundance and detection in at least three independent studies. The first [(pS)GGGAQLMPSR] was localized at S514 [37, 38]. The second [SSEVSSGSSQ(pS)PPAK] was localized at S553 [37–39]. Both were confirmed in a recent Mass Spectrometry study with high confidential values [40].

By means of site directed mutagenesis, either phosphodead exchanges to alanine or phosphomimic exchanges to glutamic acid were introduced giving four different ENP constructs with the exchanges S514A, S514E, S553A and S553E respectively (Fig. 3 F-J). We evaluated the independent transformant lines separately (Fig. 2) to obtain best information on the impacts of the mutant variants separated from possible transformation/position effects.

Assessment of GFP-signal localization in the pedigree revealed (weak) cytosolic distribution without any polar localization of GFP in both S to A single mutant constructs (ENP^S514A^; ENP^S553A^; Fig. 3F, H), whereas S to E exchanges (ENP^S514E^; ENP^S553E^; Fig. 3G, J) displayed basal (inner tissues) vs. apical (epidermis) localization of the GFP signal. In none of the lines did single S to A exchanges lead to rescue of the *pid enp* phenotype (Fig. 2). In contrast, changes from S to E always led at least to partial rescue, ENP^S514E^ performing significantly better than ENP^S553E^ (Fig. 2). Few ENP^S514E^ plants could produce (cotyledon-less) pedigree in quantities comparable to “wild-type” *EGFP-ENP* or *ENP-mGFP6* constructs (Fig. 2). ENP^S553E^ constructs could generate all flower structures but no pedigree (Fig. 2).

With these results in mind, we generated two additional constructs where both serines (at aa514 and aa553) were either replaced by alanines (ENP^S514A/S553A^) or by glutamic acids (ENP^S514E/S553E^). Interestingly, both variants resulted in perfect polarity of ENP-GFP (Fig. 3 K, L). All showed at least partial rescue (including flower organs) and 5/6 lines included plants, which produced cotyledon-less pedigree (Fig. 2). Thus, both variants ENP^S514E/S553E^ as well as ENP^S514A/S553A^ have the capability for complete rescue.

On molecular level full-length ENP (assumed to be phosphorylated at both sites ENP^S514-P/S553-P^), ENP^S514A/S553A^ (without charge) and ENP^S514E/S553E^ (with charge) should display similar although not identical characteristics such as structure, folding, association and mobility at the PM. We addressed this latter aspect with Fluorescence Recovery After Photobleaching (FRAP). Both double phospho-mimetic versions essentially displayed similar recovery dynamics as independent ENP wild-type transformants except a slightly higher recovery for the N-terminal GFP fusion (Fig. 3M). Essentially, this pattern remained stable when we altered the diameter of the region to be bleached (SFig. 8). Note, that diffusion constants and recovery times for FRAP change extremely slow with mass [41], suggesting that the behavior of the GFP-fusion construct approximates that of ENP alone.

In the next step, we tested whether a simple charge imbalance in positions S514 vs. S553 could be a cause for protein instability seen in ENP^S514A^ and ENP^S553A^, which both leave the second site free for phosphorylation. Such a situation is mimicked in the versions ENP^S514A/S553E^ and ENP^S514E/S553A^. However, these variants displayed perfect cellular polarization of ENP (Fig. 3 N, O). They were capable to partially rescue but not to generate gynoecia and pedigree (four independent transformants; Fig. 2H).

### Mutation of conserved amino acid residues often retains cellular polarity but impacts severely on functionality

The foregoing analyses demonstrated the significance of the N-terminus/central core for polarity and that of the C-terminus for function. Considering the (non-rescue) effect of the ΔN-term construct for function, we extended the analysis of the former using point mutations of highly conserved amino acids localized in the region aa1 to aa470. The proline at position 46 in the BTB/POZ domain is a conserved residue in one of two short helical regions, which form a structural turn setting a group of beta-sheets in to a (anti)parallel arrangement (Fig. 1H; SFig.1). In some non-plant BTB/POZ proteins, it is a contact site for protein interaction [26] (SFig.1). Due to its structure, proline confers a characteristic kink in the amino acid sequence of proteins. Thus, any replacement of proline should significantly alter the local protein microstructure. Plants carrying a threonine in this position (ENP^P46T^) retained perfect cellular polarity (Fig. 3P). However, only in 25% of all plants this construct led to partial rescue of the *enp pid* phenotype with bracts/cauline leaves and occasional flower structures (Fig. 3P).

Next, we replaced a highly conserved Leucine at position 144 in the linker region by aspartic acid (ENP^L144D^). In the AlphaFold prediction, L144 lies in one of a consecutive group of long and short alpha-helical structures (Fig. 1H; SFig.1). L144D led to enhanced internalization of the GFP signal. However, significant signal remained polarly localized while the *enp pid* phenotype was not rescued at all (Fig. 3Q).

We then focussed on the well conserved aa Tyrosine 409, which is part of a longer alpha-helix within a group of more or less similarly oriented helices before the start of the IDR (Fig. 1H; SFig.1). It is also part of an in-frame GLY deletion mutant (aas407-409) of the (*enp*) *mab4-1* null allele [22]. We considered both potential phosphomimic and phosphodead versions. The alteration Y409E (ENP^Y409E^) resulted in absence of any localisation, very poor presence in the cytosol and no rescue (Fig. 3R). In contrast, the alteration of Y409A (ENP^Y409A^) retained perfect polarity and achieved 10% partial rescue (i. e. only bracts/cauline leaves formed; Fig. 3S). Apparently, the replacement of highly conserved aas significantly disturbs the sequence-structural integrity of these regions, which is a precondition for functionality. However, correctly structured these regions cannot fulfil ENPs function. This is controlled by the C-terminus.

### ENP is closely associated with the PM

Confocal Laser Scanning Microscopy (CLSM) shows significant amount of ENP protein close to the PM. However, with best objectives the maximum resolution in the xy-dimension is approx. 200nm (400nm in z-dimension), which leaves significant space for a distant localization of ENP to the PM. Analysis with various algorithms (ARAMEMNON: http://aramemnon.uni-koeln.de/) does not show any prenylation or related motifs nor trans-membrane domains. However, on the basic hydrophobic (BH) scale [42] ENP displays small potential contact sites along its complete length including the C-terminus (SFig. 9). To experimentally assess potential contact between ENP and the PM we used FLIM-FRET and short (2-5min) treatments of plants with the PM-fluorophor FM4-64 (as acceptor) and GFPs from *35Sp:EGFP-ENP* and *35Sp:ENP-Cterminus-GFP6* respectively (as donors; Fig. 4A, B). The lifetime values obtained indicated close association (< 10nm) to the PM for both. Their spread towards low lifetime values (ca. 2,0 nsec) in some specimen indicated fast permeation of FM4-64 into the PM. Lifetime values with FM4-64 acceptor expressed as τ_average intensity_ were ca. 2,36 nsec (EGFP-ENP) and ca. 2,19 nsec (ENP-GFP6) as compared to controls without FM4-64, which were ca. 2,52 nsec (EGFP-ENP) and ca. 2,49 nsec (ENP-GFP6; Fig. 4 A, B).

### ENP interacts with PIN2 mainly with its C-terminus

Next, a possible FRET with the PM-integral PIN2 auxin efflux carrier was tested. PIN2 instead of PIN1 was chosen for several reasons. First, PIN1 is basally localized in the stele but as such covered by several tissue layers which aggravates the FRET analyses. In contrast, PIN2 is apically localized in epidermal and basally localized in cortex cells. PIN2 is also structurally and functionally related to PIN1, which can even replace PIN2 [43]. Additionally, PIN2 has been shown to co-precipitate with ENP/MAB4 [25].

We performed FLIM-FRET analyses with EGFP-ENP and ENP-GFP6 in combination with a PIN2-mCherry construct [44]. The latter was also combined with BRI as a negative control (Fig. 4C, D). We expected absence of interaction in this case since BRI is a PM localized brassinosteroid receptor and thus an element of a different signal transduction pathway [45]. The τ_average intensity_ life time for ENP-GFP6 alone in this experiment was ca. 2,47 nsec while it was ca. 2,33 nsec in presence of PIN2-mCherry (Fig. 4C), which gives difference of 140 psec and an energy transfer rate of E = 5,7%. This is well within the range reported for other cases [25, 46–48]. Considering the Förster distance of R_0_= 5.288 nm for the (E)GFP-mCherry pair, a distance of approx. 8.4 nm for the GFP at the ENP-C-terminus and the mCherry in the cytosolic loop of PIN2 results (this calculation assumes a *kappa^2^*orientation factor of 2/3 see SText Materials and Methods). The measurement of ENP-N-terminus vs. PIN2 exhibits only a difference of 52 psec (E = 2.1%), which is a very weak, borderline FRET (Fig. 4C). A lifetime difference of BRI-GFP alone vs. BRI-GFP combined with PIN2-mCerry was almost absent (2,52 nsec vs. 2,50 nsec). These results strongly suggest that the C-terminus of ENP interacts with PIN2 while the N-terminus is more distantly neighboured.

## DISCUSSION

ENP and MELs play an important role in auxin transport by co-operating indirectly or directly with AGC kinases, in particular PID, and PIN proteins [16, 18, 22–25]. Considering the number of aforementioned factors, which impact on the developmental effects of auxin linked to the activity of PINs, the list of (in-)direct co-operators of ENP and MELs might extend in the near future. The dissimilarity of their C-termini also suggests a corresponding number of specificities and tasks. Recently, an unexpected observation, described as haplocomplementation, fosters the view of PIN1 being part of a larger protein complex sensitive to PIN1 dosage [49]. Together with all accumulated observations this supports the existence of a PID-independent input or pathway in organogenesis with ENP as an important element.

### ENP likely contacts the PM with different parts

According to the current knowledge ENPs N-terminus and central core adopt an elongated cylindrical sphere [26–28]. ENP has no obvious lipid modification signals or transmembrane domains. At least, at its termini and its centre GFP integrations analysed in this study would have disturbed such signals. In case of the D6 protein kinase it is known that it binds polyacidic phospholipids through a lipid-rich motif [50]. According to scanning searches with the modified EMBOSS program [42] potential contact sites on a basic and hydrophobic (BH) scale are found along the entire structure of ENP. This might explain why ENP retains significant cellular polarity despite severe deletions and point mutations.

We analysed membrane association of ENP *in vivo* using FLIM-FRET. FM4-64 is a lipophilic PM stain, which initially localizes at the outer PM leaflet and is useful for studies of endocytosis [51, 52]. FM4-64 causes transient internalization of GFP-tagged PM proteins in plant cell culture cells after 10min treatments but not in the *Arabidopsis thaliana* root [52]. The treatments applied in this study indicate a FRET of ENP-GFP with FM4-64 in the outer leaflet of the PM. The internalisation of FM4-64 by endocytic processes [51, 52] cannot be fully excluded but should be marginal given the short (2-5 min) treatments. Considering the dimensions of plant PMs of approximately 6nm (hydrocarbon core and interfacial regions) [53], this is within the distance of FRET (<10nm) [54]. A possible activity of flippases [55] would transfer FM4-64 to the inner leaflet, thus bringing the dye nearer to ENP. However, FM4-64 does not appreciably flip in the PM and diffusion of FM4-64 into the cytosol could also be excluded [51]. Together, the presented FRET results show ENP being closely neighboured to the PM.

### ENPs information for tissue specific apical vs. basal cellular polarity is buried in the N-terminus and the central core

So far, it was not known whether (and if, where) ENP harboured an inherent determinant for polarity and its recognition by the cellular machinery. Our work shows, that this information is to a large part allocated in the central core of ENP. Although the BTB/POZ domain provides significant support (see ENP-ΔNterm), ENPs polarity is still realizable without this domain whereas the core is not dispensable for this (see ENP-ΔNPH3_3). The C-terminal part is largely unnecessary for polarity but supports lateral accumulation (“smile”) of polar ENP. Its interaction with PINs, such as PIN2, might likely contribute to this accumulation and backs the view of a mutual support of ENP, MELs and PINs in polarity [25]. However, PINs alone might not represent the complete machinery for ENPs polarity because ENP-ΔCterm perfectly polarizes in PIN1/2 wild-type background although the strongest interaction with PIN2 occurs with its C-terminus. Together, ENP and MEL4 (representing MELs) carry the information for polarity mainly in their central core. The read out of this information depends on the tissue where they are expressed and is valid for either apical or basal localization.

### Point mutations in the N-terminus and central core affect function rather than polarity

Partial deletions of ENP enabled us to identify the main region responsible for polarity. These deletions also lost functionality, indicating polar localization being a precondition for functionality. Considering the support of PM localized PIN activity, this was expected. Therefore, we included more subtle (point) mutations of conserved residues in our work. This showed, that the tolerance for alterations within the N-terminus and central core seems to be high with respect to polarity. All point mutation constructs except Y409E displayed significant if not perfect polarity. We are aware that the latter is not an adequate phospho-mimic since it is well known that Glutamate (and even less Aspartate) is unable to mimic either charge or the volume of pTyrosine [34]. The disturbed cellular distribution and degradation of ENP-Y409E might be caused by a severe structural impact in the helix (406D to 418E). Consequently, the “mild” exchange Y409A retains polarity and supports the view of the mentioned tolerance. Considering function, the significance of this region has to be refined as validated by rescue of the *enp pid* phenotype because all mentioned point mutations in this region at least loose functional capability. Similarly, the *inframe* deletion of G407, L408 and Y409 lead to a complete amorphic loss-of-function allele [24]. The retention of polarity in these point mutations suggests that ENPs polarity is supported by more than one (or few) highly conserved residue. This notion is corroborated by the detected (FRET) contacts with the N– and C-terminus, the lateral accumulation by addition of the C-terminus and the distribution of potential contact sites (BH scan) found along the entire ENP protein. Thus, although the C-terminus possesses the major control on functionality, a structural integrity of the whole protein is required pointing to a functional role of the protein as a whole. This would also explain why simple MEL4-ENPC-term domain swaps are insufficient to restore ENP function.

### ENPs C-terminus interacts with PINs and is an IDR whose function is critically affected by the modification of Ser^514^ and Ser^553^

Low pLDDT scores, as those given in ENPs aa471 to aa571, likely describe intrinsically disordered regions as opposed to well-defined autonomously foldable three-dimensional structures [30, 56]. For instance, a very high number of regions with low pLDDT scores of the human proteome overlaps with regions of intrinsic disorder (29, 57]. Furthermore, IDRs are not unstructured, they rather undergo disorder-to-order transitions (and vice versa) depending on special environmental and physiological conditions and take over important biological functions (33, 58, 59].

We noticed that ENPs C-terminal sequence displays an additional feature found in some IDRs. Besides numerous serines and arginines the C-terminus buries a repeating peptide motif SSSSSSRRRR (aa558-aa567). Such low-complexity regions are known in IDRs to form “collapsed globule”-assemblies as opposed to “extended coils” with alternating aa sequences [60, 61].

Finally, IDRs are prominent for harbouring, in particular serine and threonine, phosphorylation sites [34, 36], which can be linked to folding and regulatory switches [e. g. 62]. Such functional phosphorylation sites appear to be present in the C-terminus of ENP. Alterations of both tested phosphorylation sites (S514 and S553) impact on ENPs function in terms of *enp pid* rescue.

If “free” (unmodified) sites are phosphorylated by endogenous kinases, the effectivity of ENPs rescuing capability follows a pattern (Fig. 5). The pattern is, that the stronger the bias between S514 and S553, charge vs. no charge and charge of glutamic acid vs. that of phosphorylated serine, the lower the rescue success (Fig. 5; Fig. 2H, J). The single phosphodead constructs are unstable either because phosphorylation at the “free” second site is impossible and/or causes instability. This problem is absent in the next constructs, which have a fixed (modified) translatable sequence. Here a bias in charge is less favorable than a mix of negative charge provided by phosphomimic and endogenous phosphorylation. These in turn are less effective than identical modifications on both sites even when both are phosphodead. The wildtype constructs undergo endogenous phosphorylation at both sites and perform best. The fused GFP proteins might cause slight differences to the *pid* single mutant with respect to (residual) flower formation and fertility. Considering the shown interaction of the C-terminus with PIN2 it is likely, that biased and non-biased modifications respectively might be structurally detrimental or sub-optimal for this interaction as compared to full wild-type phosphorylation. This awaits in depth structural analyses. The instability of the single phosphodead constructs could indicate an interesting side effect. The detrimental effects of incompletely phosphorylated ENP guarantees kinase activity until phosphorylation is completed – kind of safeguard or counting mechanism. Taking together, the C-terminus of ENP likely represents an intrinsic disordered region, which is essential for ENPs activity and can be modulated by modification of selected target serines. This is at least partly attributed to a PIN-interaction, whose strength depends on its phosphorylation status.

The accumulated data of this and previous studies delineate a model of how ENP independently impacts on organogenesis by supporting PIN polarity (Fig. 6). PID activity has a major impact on PIN1 polarity and activity in *enp* mutant background (Fig. 6A, B). ENPs contribution is then visible by two effects. On the molecular level, the abundance of PIN1 carriers is reduced at the PM [22]. This has mild but detectable consequences on the developmental level as seen by fused sepal organs [16]. In the *pid*-background (Fig. 6C), residual (polar) PIN1 maintains low auxin flux [16]. This is enabled by the interaction of PINs and ENPs (phosphorylated) C-terminus and results in plants with (partly) pin-formed stems and stems with abnormal, fertile flowers (this study). In old *pid* embryos the PIN1 population is distributed on apical, lateral as well as basal regions of the PM [16]. In *enp pid* double mutants (Fig. 6D), the cellular PIN1 population has completely shifted to lateral and basal PM regions and all plants only develop blind stems [16]. In this study, the ENP-ΔCterm construct mimicks the original *enp* allele and suggests that the lateral/basal shift of PIN1 is due to the absence of ENPs C-terminus while ENP remains apically localized. This model implies that *pin pid* [17] resemble *enp pid* double mutants. Some of the aforementioned factors, whose mutants result in cotyledon– and flower-less phenotypes in the *pid* background, might also contribute to the PID-independent input.

## Materials and Methods

### Plant material, growth conditions and seedling culture

*Arabidopsis thaliana* (ecotype L*er*-0), EMS-induced single/double mutants and transgenic construct lines were grown according to conventional procedures under continuous light or 12 hrs light/12 hrs dark cycles (for details see SText Materials and Methods).

### Cloning and site directed mutagenesis, deletion and domain swap constructs

Briefly, *ENP* and *MEL4* Wild-type full-length cDNA clones (pda08292 and pda10515, Riken Bio Resource Center, Japan) were used as starting material for further cloning by conventional restriction-ligation or Gateway technology (Thermo Fisher Sc.). For deletion, domain swap and site directed mutagenesis constructs, appropriate primers extended with restriction sites recombination sites were used (see SText Materials and Methods). For site directed mutagenesis the Quick Change II (Agilent) or the Q5 Site Directed mutagenesis Kit (NEB) according to the supplieŕs instructions were used.

### Sequencing

We assessed critical regions on all levels of cloning and (after) transformation in *E. coli*, *A. tumefaciens* and *A. thaliana* with appropriate primers by sequencing (EUROFINS sequencing services).

### Plant transformation

Plants were transformed according to conventional methods using *Agrobacterium tumefaciens* strain GV3101.

### Chemicals and pharmacological studies

Seedlings were treated with 10mM PBA (MERK) as described [31] and with FM4-64 (1,7µM-2µM; ThermoFisher Sc.) for 2-5 min, washed in water and processed for Imaging and/or FLIM-FRET analysis.

### Immunocytochemistry

PIN1 localization in embryos transgenic for 35Sp:EGFP-ENP used PIN1 primary rabbit antibody incubation (1:1000; 4h, 37°C) and secondary rabbit Cy3-Antibodies (BSA/PBS for 3.5h at 37°C; Jackson ImmunoResearch/USA supplied by Dianova/Hamburg). After repeated washes with PBS and H_2_0 the embryos were embedded in Citifluor antifadent mounting medium and covered with a coverslip, stored at 4°C or –20°C or immediately processed for imaging.

### In situ hybridization

Verification of mRNA patterns in embryos were as previously described [4, 16].

### Confocal Laser Imaging Microscopy (CLSM) and FRAP analysis

Imaging was done with Olympus FV1000 or FV3000 and 20X/0.75 NA air Plan-Apochromat or 63X/1.2 NA Plan-Apochromat water objective and TCS SP8 Leica equipped with a 63XW/NA 1.2 Plan-Apochromat water objective using the corresponding company software. Imaging used excitation laser lines 488nm Argon, 488nm diode, 515nm diode or 561nm diode lasers and appropriate detection (windows) Olympus PMT or GAsP detectors or Leica TCS SP8 HyD or PMT detectors. While HighVoltage detector setting was adjusted according to signal strength, non-linear signal amplification was not performed. Also, other than zero threshold setting (“Offset”) was regularly avoided.

FRAPs were performed with the TCS SP8 CLSM with 20µm (and 40µm) bleach spot diameters. After bleaching at high intensity with the 488nm Argon laser, fluorescence intensities of the same and unbleached control regions (same spot size) were measured at different time intervals and normalized according to I_n_ = (I_t_-I_0_)/(I_I_ – I_0_), where I_t_ is the value of the recovered fluorescence intensity at any time t, I_0_ is the first post-bleach fluorescence intensity and I_I_ is the initial (pre-bleach) fluorescence intensity.

### Measurement of polar GFP-signal distribution (“smile” analysis)

Cells within the epithelial and cortex region were measured if at least five were suitable for measurements. For each cell three different measurements were made. The first of the apical membrane, the second of the residual membrane and the third measurement was of the whole length of the GFP signal (SFig. 8). Then GFP-signal lengths over apical membrane lengths, and GFP-signal length over total cell circumference (= apical + residual membrane length) were calculated and subjected to individual *t-*Tests.

### Fluorescence Lifetime Imaging Microscopy (FLIM) and Förster Resonance Energy Transfer (FRET) measurement

The lifetimes (τ) of donor Fluorophores (EGFP, GFP6) without and after non-radiative energy transfer to acceptor molecules (FM4-64, mCherry) were measured with the aid of a PicoQuant-Kit for Time Correlated Single Photon Counting (TCSPC) connected to a FV3000 Olympus CLSM following PicoQuant instructions. Energy transfer efficiency (E) was calculated according to: E = 1 – τ_DA_/ τ_D_, where τ**_DA_** is the fluorescence lifetime of the donor in presence of the acceptor and τ**_D_** is the fluorescence lifetime of the donor alone. For approximation of the distance (r) between (E)GFP and mCherry pairs we took R_0_ (the Förster distance at 50% energy transfer) from the “FPbase FRET Calculator (at https://www.fpbase.org/fret/) and used the equation for E expressed as E = R_0_^6^/(r^6^+R_0_^6^) [54].

For details to all listed Materials and Methods see SText Materials and Methods.

## Acknowledgements

We thank T. Sieberer, M. Nakamura, U. Mayer, G. Jürgens and the Nottingham Arabidopsis Stock Centre (NASC) for construct lines BRI-GFP, PIN2-mCherry, PIN1-Antibody and *Arabidopsis* lines and O. Peis for technical support. We gratefully acknowledge financial support by: the Deutsche Forschungs-Gemeinschaft in the initial phase of the project (DFG To134/8), a fellowship of the Bayerische Eliteförderung (M. L.), a TUM Laura Bassi equal opportunity program fellowship (M. S. M) and TUM School of LifeSciences Studienkoordination supports of FP, BSc and MSc projects (U.B., B. S., M.S.L. and N.Y). We are indebted to A. Gierl for consistent support and K. H. Schneitz for hospitality. Microscopic analyses were carried out with equipment from the Center of Advanced Light Microscopy (CALM) of the School of Life Sciences, Technische Universität München.

## Author contributions

M.S.M. and R.A.T.R. designed research, M.S.M., N.Y., M. L., U.B., B.S.F., B.S., M.S.L. and R.A.T.R. performed research, R.A.T.R. wrote the paper.

## Competing interests

The authors declare no competing interest.

## Notes

### Competing Interest Statement

The authors have declared no competing interest.

### Summary of Updates

The manuscript has been extended by updating Supplemental files including 1. Supporting Text: Materials and Methods and 2. Supplementary figures and their legends.

