## Supporting Text for "Separate domains of the *Arabidopsis* ENHANCER OF PINOID drive its own polarization and recruit PIN1 to the plasma membrane"

### Supporting Text: Materials and Methods

#### *Plant material, growth conditions and seedling culture*

*Arabidopsis thaliana* (ecotype L*er*-0) and EMS-induced single and double mutants (*enp pid15/enp pid-15 = laterne*; *enp pid-15/enp* +; *pid-15/pid-15* as described [16] and transgenic construct lines were surface sterilized (3x wash with $H_2O$, 1x 3-4% bleach for 20 min, 3x $H_2O$ wash) and sown on 1/2 MS selection media without/with appropriate antibiotic (i. e. Hygromycin 13mg/l; Phosphinotricin 25mg/l), stratified for 1-2 days at 4ºC in the dark and grown for 5 to 14 days at constant illumination (Sanyo light chamber with 19±2Cº). For propagation of lines, seeds were then transferred directly sown on soil under continuous light as described [16] or 12hrs light/12 hrs dark and 21ºC ±3º, between 100-200 [µMol $m^{-2}$ $sec^{-1}$] and ca. 50% humidity. We did not observe *enp pid-15/enp pid-15* (*laterne)* phenotype alterations under these growth conditions.

#### *Cloning and site directed mutagenesis, deletion and domain swap constructs*

The ENP Wild-type full-length cDNA clone pda08292 (RAFL09-39-B04; Riken Bio Resource Center, Japan) was used as starting material for further cloning of ENP constructs. The ENP coding sequence was amplified (Proofstart Taq Polymerase, Quiagen) and blunt ligated into SmaI-digested pBluescriptII KS+ (Stratagene). Using these plasmids, ENP was amplified with primers homologous for 5´and 3´-end extended by EcoRI and BamHI restriction sites (see below) and cloned via EcoRI/BamHI sites into the pEGAD vector [63] to give *35Sp:EGFP-ENP*. A second construct lacking the first two introns (amino acid no.1 to 53; SFig. 1) was generated in an analogous way to give *35Sp:EGFP-ENP-*$\Delta$*Nterm* (SFig. 2).

All other *ENP* constructs were generated using GATEWAY technology (Invitrogen/Thermo Fisher Scientific). *ENP* full length with flanking attB1/B2 sites was

PCR amplified using primers overlapping with the corresponding 5´-/3´-ends and extended with attB1/B2 sites for recombinant cloning.

Recombination of the amplified fragment with vectors pDONR207 and pDONR221 using BP$^{TM}$ clonase gave full length *ENP* ENTRY clones.

Site directed mutagenesis constructs were generated with full length *ENP pDONR207/R221* ENTRY clones as starting material using the Quick Change II (Agilent) or the Q5 Site Directed mutagenesis Kit (NEB) according to the supplier´s instructions using appropriate primers (see below). For constructs with ENP$^{S514AS553E}$ and ENP$^{S514ES553A}$ the existing construct with ENP$^{S514ES553E}$ was taken for a further site directed mutagenesis round.

Deletion constructs (except *35S:EGFP-ENP-∆Nterm*) were generated using full length ENP clones as template for PCR amplification with 5´and 3´ attB1/B2 flanked primers for the full length clone combined with primers starting at the corresponding start/end position in *ENP* and also extended with attB1/B2 sites (primers see below). Finally, attB1/B 2 flanked fragments were recombined with BP$^{TM}$ clonase to give deletion ENTRY clones.

Full length, point mutagenized and deletion ENP ENTRY clones were recombined with pMDC83 binary vector [64] using LR$^{TM}$ clonase. In these clones *ENP* precedes *GFP6his*. For instance, full length *ENP* with C-terminally fused GFP6 (as opposed to *35Sp:EGFP-ENP* with N-terminally fused EGFP*)* gave the DESTINATION construct *35Sp:ENP-GFP6*.

For generation of a full length *MEL4/NPY4* construct, the coding sequence of the cDNA clone pda10515 (RAFL17-11-E06; Riken Bio Resource Center, Japan) was taken and cloned into pDONR207 using GATEWAY technology in the same way as for *ENP*. The *MEL4*-ENTRY construct was recombined with the pEarlyGate102 vector to give *35Sp:MEL4-EYFP* (for primers see below).

Two *MEL4-ENPCterm*-domain swap constructs were generated by extended overlap PCR and via ENTRY clones finally inserted into pMDC83 binary vector [64] using LR$^{TM}$ clonase (see primers below).

For construction of *35Sp:MEL4-ENPCterm_long-GFP6*, *MEL4* (Riken clone pda10515) was amplified with primers *MEL4_att1FW* and *MEL4_ENPCterm_long*. The resulting fragment includes an att1-site followed by the *MEL4* 5´-end until base 1356 (including the MEL4 aa 452) and finally an extension of 30bp starting from ENP base 1411 to 1441(aa 471 to 481). The Cterm-end of ENP starting at bp1411 to the end at 1713 (aa 571) was amplified with *ENPCtermLong FW* and *ENP_att2REV.* The fragments were melted, hybridized and PCR-amplified to give an att1/att2-flanked fusion of MEL4 from amino acid 1 to 452 and ENP amino acid 471 to 571.

The construction of *35Sp:MEL4-ENPCterm_short-GFP6* was performed in the same way using *MEL4_att1FW* and *MEL4_ENPCterm_short* for the MEL4 part (amino acid 1 to 481) and *ENPCtermShort FW* and *ENP_att2REV* for the ENP part (amino acid 500 to 571). The fragments were melted, hybridized and PCR-amplified to give an att1/att2-flanked fusion of MEL4 from amino acid 1 to 481 and ENP amino acid 500 to 571.

*Primers for ENP cloning into pEGAD*

(EcoRI and BamHI sites underlined).

*For 35Sp:EGFP-ENP:*

5´- CCT<u>GAATTC</u>ATGAAGTTCATGAAGCTAGGGTCTA -3`

5´- AC<u>GGATCC</u>TCACGATATCGAATGTCTG -3´

*For 35Sp:EGFP-ENP-∆Nterm:*

5´- GCT<u>GAATTC</u>ATGCAGCGACTGGTTTTT -3´

5´- GAC<u>GGATCC</u>TCACGATATCGAATGTCT -3´

*Primers for ENP GATEWAY cloning*

*For 35Sp:ENP-$GFP_6$ construct:*

*ENP_att1FW*

5´- GGGGACAAGTTTGTACAAAAAAGCAGGCTTCATGAAGTTCATGAAGCTAGGG TC -3´

*ENP_att2REV*

5´- GGGGACCACTTTGTACAAGAAAGCTGGGTCCGATATCGAATGTCTGCGGCG -3´

*35Sp:ENP-ΔCterm-$GFP_6$* construct (for 5´end see full length *ENP*):

5´- GGGGACCACTTTGTACAAGAAAGCTGGGTTGAGCTGCTCAAAGTAGAGAA CTTG -3´

*35Sp:ENP-ΔNPH3_3-$GFP_6$* construct (for 5´end see full length *ENP*):

*5´*- GGGGACCACTTTGTACAAGAAAGCTGGGTTCAACAAAATTCCATTTCCTAA AAC -3´

*35Sp:ENP-CtermOnly-$GFP_6$ construct* (for 5´end see full length *ENP*):

5´- GGGGACAAGTTTGTACAAAAAAGCAGGCTTCATGCACAGCCCCGTGGC GTCT -3´

*Primers for site directed mutagenesis (mutated triple underlined)*

Site directed mutagenesis was performed with kits from Agilent or New England Biolabs respectively according to the supplier´s instructions.

Primers used in the Qick Change II procedure (Agilent)

*ENP_P46T*

5´- CACCTCCATAAGTTCACGCTGCTATCGAAGAGC -3´

5´- GCTCTTCGATAGCAGCGTGAACTTATGGAGGTG -3´

*ENP_L144D*

5´- GGAAAGACTCAATCATTGTGGATCAGACAACAAGATCTCTTC -3´

5´- GAAGAGATCTTGTTGTCTGATCCACAATGATTGAGTCTTTCC -3´

*ENP_Y409E*

5´- CCGATACACGACGGTCTCGAGAAAGCCATTGACACT -3´

5´- AGTGTCAATGGCTTTCTCGAGACCGTCGTGTATCGG -3´

*ENP_Y409A*

5´- CCGATACACGACGGTCTCGCTAAAGCCATTGACACTTTCATG -3´

5´- CATGAAAGTGTCAATGGCTTTAGCGAGACCGTCGTGTATCGG -3´

*ENP_S514E*

5´- GAAGCAAGAGCACGAGGGAGGGTGGTGGTGCACAGCT -3´

5´- AGCTGTGCACCACCACCCTCCCTCGTGCTCTTGCTTC -3´

*ENP_S514A*

5´- GCAAGAGCACGAGGGCTGGTGGTGGTGCAC -3´

5´- GTGCACCACCACCAGCCCTCGTGCTCTTGC -3´

*ENP_S553E*

5´- TCTGAGGTTTCTTCTGGAAGCTCACAAGAGCCGCCAGCCAAGTC -3´

5´- GACTTGGCTGGCGGCTCTTGTGAGCTTCCAGAAGAAACCTCAGA -3´

*ENP_S553A*

5´- AGGTTTCTTCTGGAAGCTCACAAGCTCCGCCAGCCAA -3´

5´- TTGGCTGGCGGAGCTTGTGAGCTTCCAGAAGAAACCT -3´

*Primers used in the Q5 site directed mutagenesis kit (NEB)*

*ENP(S514E) to E514A*

5´- GAGCACGAGGGCTGGTGGTGGTG -3´

5´- TTGCTTCCTCTGCTTTTCTTG -3´

*ENP(S553E) to E553A*

5´- AAGCTCACAAGCTCCGCCAGCCA -3´

5´- CCAGAAGAAACCTCAGAGC -3´

*Primers for MEL4/NPY4 and MEL4/ENP domain swap constructs.*

For *35Sp:MEL4-EYFP:*

*MEL4_att1FW*

*5´-* GGGGACAAGTTTGTACAAAAAAGCAGGCTTCATGAAGTTTATGAAACT

TGGAA -3´

*MEL4_att2REV*

5´- GGGGACCACTTTGTACAAGAAAGCTGGGTCAAACTCTTTCTCATGGTC

CCATT -3´

*MEL4_ENPCterm_long*

5´- CGAAGCCGCAACAGACGCCACGGGGCTGTGATTGGCTCTAATCTGTTC

AAAGAAGAGAAC -3´

*MEL4_ENPCterm_short*

5´- GCTTCCTCTGCTTTTCTTGCTCAGCTCCACAAACTCTTTCTCATGGTCCC

ATTCATCATCCTC -3´

*ENPCtermLong FW*

5´- CACAGCCCCGTGGCGTCTGTTGCGGCTTCGTCACACTCGCCGGTTGAG

AAG -3´

*ENPCtermShort FW*

5´- GTGGAGCTGAGCAAGAAAAGCAGAGGAAGCAAGAGCACGAGGAGTGGT -3´

***Sequencing***

We assessed critical regions (point mutations, deletions, swaps) on all levels of cloning and (after) transformation in *E. coli*, *A. tumefaciens* and *A. thaliana* with appropriate primers (SIText) by sequencing (EUROFINS sequencing services).

Initially as comparison to the genetic assessment of endogenous *ENP/enp-1* and *PID/pid-15* respectively, *enp pid/enp pid (laterne*) seedlings with transgenic *35Sp:EGFP-ENP* were genotyped by means of pyrosequencing with primers as previously described [16] (SText).

*Primers for assessment of constructs*

Sequencing was performed with conventional insert flanking primers for cloning vectors, sometimes with primers for cloning (see above) and the following primers (in particular for pMDC83 constructs).

*ENPCterm_FW*

5´-CAG AAC GAG AGA CTT CCA CTA-3´

*GFP_to_ENP_rev*

CCT TCA CCC TCT CCA CTG ACA G

*ENP_NPH3_1_FW*

GTT GCA AGG TGG TTA CCA GAA

*35S_ENP_FW*

CAC TGA CGT AAG GGA TGA CGC A

*35SLinker_ENP_FW*

ACA GCG ACA GCT ATC AGT TGC

*ENPLinkerREV*

ATC TTT CGG AAT CAC ATT CTC

*Pyrosequencing Primers*

*Target: enp-1 allel:*

Biotin-5´- GCACGCTGCACAGAACGA-3´

5´-CGGGGCTGTGATTTG-3´

5`-GCCACGGGGCTGTGATTT-3`.

*Target pid-15 allel:*

Biotin-5'-CTTGACGACGGAAGAAGGAATC-3'

5'-CATGCGCGGAATTTGATTT-3'

5'-GATCCGACTAAAAGACTTG-3.

***Plant transformation***

*Agrobacterium tumefaciens* strain GV3101 [65] was used to transform *Arabidopsis thaliana* plants (either wild-type L*er*-0 and Col-0 or *enp pid/enp* + always L*er*-0).

The „floral dip"-Method [66] was followed for plant transformation.

***Chemicals and pharmacological studies***

Chemicals were purchased from MERC (Phenylboronic acid, PBA) and ThermoFisherScientific (FM4-64). PBA (10mM) was applied as described [31] to seedlings carrying *35Sp:MEL4-EYFP* and then directly mounted on slides with the same solution for CLSM analysis.

We experienced and considered that infiltration with FM4-64 can be variable between individual seedlings. Therefore, seedlings were stained with FM4-64 (1,7µM or 2µM) for 2-5 min, washed twice in water, one wash for 5-15 min and a second wash for 1min. The seedlings were then processed for Imaging and FLIM-FRET analysis.

***Immunocytochemistry***

Fixation and wash on 1st day embraced the following steps: ovules with embryos (heart to torpedo stage) were isolated from the siliques and incubated for 1h under vacuum in 1ml 4% PFA in MTSB (50mM PIPES; 5mM EGTA; 5mM $MgSO_4$; pH7) with 10µl 10% Triton and then either further processed or stored at 4ºC. Ovules were then washed 4 x 10 min with MTSB/ 0,1% Triton, 2 x 10 min with PBS (137mM NaCl; 2,7mM KCl; 10mM $NaH_2PO_4$; 2mM $KH_2PO_4$; pH7)/ 0,01% Triton, 2 x 5 min with PBS (on ice) and finally 1 x 5 min with $H_20$. Then ovules were placed with as less liquid as possible on gelatinized microscope slides, covered with 22x22mm coverslips and gently squeezed to press out the embryos from the ovules. Excess liquid was aspirated with Kleenex, the slides were submerged into liquid nitrogen and the coverslip blown up with a razor blade. The slide was dried o/N at RT (if necessary stored at -20ºC).

2nd day: Specimen on the slide were surrounded with a Pap pen and covered with 1ml MSTB for 10min. Afterwards MSTB was removed and the specimen covered with 200ml 2% Driselase in MSTB and incubated for 25-45 min (depending on the Driselase batch). Subsequent wash was 4 x 5 min with PBS (1-1,5 ml per slide). Permeabilization of membranes was achieved with 200µl 10% DMSO/3% NP40 in MTSB and incubation for 1h at RT followed by 6x5 min wash in PBS and incubation with 3-5% BSA in PBS for 1h at 37ºC (optional o/N at 4ºC). Then incubation followed with 200µl PIN1-AB (1:1000) for 4h at 37ºC or o/N at 4ºC (in a humid chamber sealed with parafilm).

3rd day: wash steps 3 x 10 min with PBS/ 0,01% Triton and 3 x 10 min with PBS were followed by incubation with 200 µl Cy3 secondary-AB (1:600) in BSA/PBS in a sealed humid chamber for 3.5hrs at 37ºC. After washing 4 x 10 min with PBS and 2 x 10 min with $H_20$ the embryos were embedded in 300 µl Citifluor antifadent mounting medium and covered with a coverslip. The embryos were then analyzed under CLSM or stored at -20ºC or at 4ºC for several months until imaged.

***In situ hybridization***

On the first day, fresh plant material/siliques were fixed in 4% Paraformaldehyde in PBS (137mM NaCl; 2,7mM KCl; 10mM $NaH_2PO_4$; 2mM $KH_2PO_4$; pH7). Approx. 15-20 siliques were partly cut with a scalpel, submerged in 1.5ml solution on ice, repeatedly vacuum aspirated and carefully ventilated every 15min. Then the fixing agent was replaced and the material stored o/N at 4ºC.

On the next day, the fixing agent was washed twice with PBS in $H_2O_{bd}$ for 30min and the PBS was then progressively replaced by increasing concentrations of EtOH in 0.85% NaCl (w/v) under continuous shaking for 1h at 4ºC, i. e. with 30%, 40%, 50%, 60%, 70%, 85% and finally $\geq$95% EtOH and stored o/N at 4ºC. Interruption at 70% EtOH was possible for storage at 4ºC for up to four months.

All the next steps were performed at room temperature (RT; Eppendorf Thermomixer, continuous shaking). The EtOH led to the extraction of chlorophyl while the addition of eosin should enhance the contrast of the specimen in the wax. Plant material was incubated twice in new 100% EtOH/0.1% eosin (w/v) for 30min. The next steps were: 2x incubation for 1h in 100%EtOH, then incubations in Histoclear/EtOH (1:3, 1:1, 3.1 each 1h) and finally 2x 1h incubation in 100% Histoclear. The material was then incubated without shaking in 100% Histoclear (Plano) mixed with 3-4 Paraplast chips o/N followed by 42ºC incubation in the Thermomixer with additional Paraplast chips for 30min.The tubes were transfered to a 60ºC oven and the solution continuously saturated by replacing 50% of the solution with molten Paraplast until reaching pure Paraplast (took at least 4hrs), which was replaced another time at 60ºC was o/N. In the next three days the Paraplast was replaced by fresh one 2-3 times a day. Then the tubes were placed at RT until complete hardening.

The resulting blocks were eventually trimmed and thin sections (7-8µM) were produced with a microtome (Reichert&Jung), placed on water drops on SuperFRost Ultra Plus slides (Menzel), dried at 42ºC on a heating plate and either stored at 4ºC or further processed.

Single stranded digoxigenin-labelled RNA probes were generated from cloned genes of interest. After linearization with appropriate restriction enzymes (not producing 3´overhangs) and purification through GFX$^{TM}$ columns, 1µg of plasmid template was transcribed with either T7, T3 or SP6 RNA polymerase, depending on the promoter and orientation in the vector according to the supplier instructions (1x Transcription buffer, 1x Digoxigenin labelling mix with DIG-UTP, 20U RNase inhibitor, 40U RNA-Polymerase in $H_2O_{DEPC}$, Roche). Hybridization probes were stored in $H_2O_{DEPC}$ and 20U RNase inhibitor at -20ºC and alkali hydrolyzed to fragments of ca. 150 nts length (hydrolyzation buffer: 200µl 0.5M $Na_2CO_3$, 160µl 0.5M $NaHCO_3$, 600µl $H_2O_{DEPC}$). The fragments were EtOH precipitated, washed with 70% EtOH, resolved in 50% deionized formamide and stored at -20ºC.

Before use, the slides were further processed either in glass or polyoxymethylene racks. Then next treatments included: 2 x 10min wash in 100% Histoclear, 2x 2min in 100% EtOH, 1min in 95% EtOH and $H_2O_{bidest}$, 1min in 90% EtOH and $H_2O_{bidest}$, 1min in 80% EtOH and $H_2O_{bidest}$ and 0.85% NaCl, 1min in 60% EtOH and $H_2O_{bidest}$ and 0.85% NaCl, 1min in 30% EtOH and $H_2O_{bidest}$ and 0.85% NaCl, 2min in 0.85% NaCl followed by 2x wash in PBS. In the next step, the sections were Proteinase K digested (1µg/ml conc. in Proteinase K buffer: 100mM Tris pH7.5, 50mM EDTA pH8.0) for 20min and the reaction stopped by incubating the slides in PBS/0.2% Glycine followed by 2x PBS wash. The subsequent post-fixation was for 2 min in 4% PFA in PBS followed by 2x PBS wash.

Then the slides were incubated in 0.6 ml acetic acid anhydride mixed with 200ml Triethanolamin (TEA) solution (2,68ml TEA in 200ml $H_2O_{bidest}$) for 2 min, washed 2x 5min in PBS and then incubated in 0.85% NaCl for additional 5 min. A dehydration series followed with the same steps from smallest to the highest (100%) EtOH concentration. Hybridization was performed in a wet chamber at 55ºC in hybridization buffer (0.3M NaCl, 10mM Tris pH7.5, 1mM EDTA pH8.0, 50% deionized formamide, 10% dextransulfate, 1x Denhardts, 05mg/ml tRNA in $H_2O_{bidest}$) with probe (40ng/1kb probe length and 100µl hybridization buffer/slide). The probe buffer mixture had been denatured for 2min at 80ºC and immediately placed on ice. The sections were incubated at 45-60ºC (depending on the probe) o/N under coverslips in the wet chamber. The next day slides were washed for few minutes at 55ºC with 2x SSC (0.3M NaCl, 20mM $Na_3$-Citrate·$2H_2O$) until coverslips fell off. Then 4x wash in 0.2x SSC at 55ºC for 20min were followed by 10min wash at 37ºC in 0.2x SSC and incubation for few minutes in PBS. The slides were twice covered with 1-2 ml blocking solution (Roche 10% blocking reaction solution in maleic acid buffer, 100mM maleic acid pH7.5; 150mM NaCl) per slide for 30min and followed by 45min wash in 2ml BSA wash-buffer (1% BSA, 0.3& Triton-X 100, 100mM Tris-HCl pH7.5, 150mM NaCl in $H_2O_{bidest}$) all at RT and in the wet chamber. After discarding the wash-buffer, the sections were covered with120µl of the antibody solution (Anti-Digoxigenin-AP Fab in BSA wash buffer 1:1250) and a cover slip and incubated for 1h 30min at RT in the wet chamber. Then two times coverslips and buffer were discarded, slides immediately covered with fresh 1-2ml wash buffer and incubation continued. Afterwards, the buffer was discarded and the slides washed and incubated in small boxes with TNM-50 buffer (8 µl NBT/BCIP [Roche] in 1ml TNM-50: 100mM Tris pH9.5, 100mM NaCl, 50mM $MgCl_2$) for 2x15min with careful shaking at RT. Then slides were covered with 120µl staining solution and incubated in a wet chamber (paper saturated with TNM-50) o/N in the dark at RT for 12hrs or longer. The staining reaction was stopped in TE for 5 min, the TE discarded, the slides carefully dried, covered with the mounting medium Entellan

(Merck) and a coverslip. After hardening of the Entellan the sections were inspected under a Zeiss Axiophot.

***Confocal Laser Imaging Microscopy (CLSM)***

Images were taken on an Olympus FV1000 or FV3000 with a 20X/0.5 NA or 20X/0.75 NA Plan-Apochromat air objective or 63X/1.2 NA Plan-Apochromat water objective using the corresponding Olympus FV-10/FV-31 software.

(E)GFP was imaged using a 488nm Argon laser (FV1000) or 488nm diode laser line (FV3000) for excitation and spectral detection with Olympus PMT or GAsP detectors between 495nm and 550nm.

EYFP was imaged using a 515nm Argon (FV1000) or diode laser line (FV3000) for excitation and spectral detection with Olympus PMT or GAsP detectors between 523nm and 600nm (CHECK). Alternatively, EYFP can be reasonably visualized as well with a 488nm Argon laser (FV1000) or 488nm diode laser line (FV3000) for excitation and spectral detection with Olympus PMT or GAsP detectors between 495nm and 550nm.

FM4-64 and mCherry were imaged using or 561nm diode laser line (FV1000, FV3000) for excitation and spectral detection with Olympus PMT or GAsP detectors between 580nm and 650nm.

While HighVoltage(Olympus)/Gain(Leica) setting was adjusted according to signal strength, non-linear signal amplification (called “Gain-setting” in Olympus microscopes) was not performed. For image representations, threshold setting (“Offset“) was mostly set to zero. For all specimen subjected to FLIM it was always zero.

For post-acquisition inspection/analysis Olympus FV-10/FV-31, FIJI/ImageJ or OMERO software was used. For visualization (E)GFP was false-colored using a linear green look-up-table, EYFP was false-colored using a linear yellow look-up-table and FM4-64 was false-colored using a linear red/magenta look-up-table. Images were assembled using Adobe Photoshop with adjustments no other than brightness and contrast. In case of the representation of weak GFP-fluorescence signals, in particular for the constructs *35Sp-ENP$^{S514A}$-GFP6* and *35Sp-ENP$^{S553A}$-GFP6,* brightness and contrast had to be significantly elevated.

***Measurement of polar GFP-signal distribution ("smile" analysis)***

The extension of GFP-signal for ENP-GFP6 vs. ENP-ΔCterm-GFP6 constructs in *Arabidopsis thaliana* was compared. For ENP-GFP6 184 cells in images of 21 different seedlings whereas for ENP-ΔCterm-GFP6 107 cells in 14 seedlings were measured. Only cells within the epithelial and cortex region were measured and only images were chosen where at least five cells were suitable for measurements. For each cell three different measurements were made. The first for the apical membrane, the second of the residual membrane and the third measurement was of the whole length of the GFP signal (SFig. 8).

In the next step a ratio for GFP-signal length over apical membrane length, and GFP-signal length over total cell circumference (= apical + residual membrane length) was calculated for each cell. The results were now percentages of GFP signal compared to membrane length to quantify and compare membrane localization of ENP-GFP6 and ENPΔCterm-GFP6. Between 5-10 cells were measured the averages for the ratio of GFP-signal and membrane were calculated for each individual seedling to ensure that the impact of a single seedling on further analysis was always the same. Following these calculations, boxplots were created out of the average ratios per seedling and individual t-Tests assuming unequal variance (Welch´s t-Test) were carried out. Both, one-tailed and two-tailed tests gave $p<0.0001$. One for GFP-signal length over total cell circumference and the other for GFP-signal length over apical membrane length.

#### *Analysis of protein mobility by Fluorescence Recovery After Photobleaching (FRAP)*

FRAP is to support the understanding of the mobility of proteins by diffusion, exchange and binding, which impact on the recovery of fluorescence after bleaching. This technique supports the understanding of the complex nature of the interactions for instance between membrane associated or integrated proteins in a lipid environment. In the presented study, FRAP was applied solely to to monitor possible differences between independent constructs in particular wild-type and constructs with double mutations in S514 and S553. All constructs were analyzed in L*er*-0 ecotype genetic background.

We used a TCS SP8 Leica CLSM equipped with a 63XW/NA 1.2 PlanApochromat water objective and a 488 Argon-Laser line. The settings for imaging were: 1% (0.8 µW) - up to 10% laser power (of initial 20% Argon laser power setting). For FRAP analyses, scanning was adjusted to 256X256 or 512X512 pixel. The bleach was applied at 80% -100% 488nm Argon laser power on an area of ca. 20µm$^2$. A bleaching experiment was at least repeated three times for every construct. Additional FRAPS were performed with altered bleach spot sizes (SFig. 8).

The so called “Fly-mode“ (Leica corp.) was applied. Here, the scanning beam bleaches during the forward sweep with intensity modulation for bleaching in the ROI and records the signal with scanning intensity during the fly-back. This process ensures a near-instantaneous recording of the post-bleach intensity for every bleached line.

The comparisons with unbleached regions and control regions without fluorescence and experiments also revealed the known robustness of GFP against weak laser intensity. Thus, GFP can be well photobleached with high intensity confocal laser pulses and imaged without significant photobleaching using low-intensity illumination [67-69 and references therein].

Before bleaching 3-5 prebleach images at time increments of 0.65 sec (alternatively 0.5-2.5 sec) were taken, which were then followed by the bleach (three repetitions at 0.65 sec). After bleaching scanning intervals for the first vs. later frames were changed, i. e. extended in the latter to adjust for larger intensity increments during the first seconds of recovery. Up to 150 post-bleach measurements (depending on the construct analyzed) were taken. The time increments of post-bleach images were 0.65sec for the first 40 images and longer for the following images (e. g. 1sec for the next 30 images and 5sec for the following). The fluorescence intensity data were normalized according to:

$$I_n = (I_t - I_0)/(I_I - I_0)$$

where $I_t$ is the value of the recovered fluorescence intensity at any time t (mean of x-independent measurements), $I_0$ is the first post-bleach fluorescence intensity (mean of x-independent measurements) and $I_I$ is the initial (pre-bleach) fluorescence intensity (mean of x-independent measurements) [70]. The end-value(s) $I_n$ of a measurement converge to a threshold that indicates the mobile and immobile fractions $F_m$ and $F_i$ respectively. The latter can be determined as $F_i = 1 - F_m$.

Care was taken to monitor movements of the seedling along the three axes and was monitored by measurement of a non-bleached control membrane. We also selected a particular region for these measurements, the meristematic region where the epidermis becomes the outer tissue.

***Fluorescence Lifetime Imaging Microscopy (FLIM) and Förster Resonance Energy Transfer (FRET) measurement***

*General considerations*

In the process of Förster Resonance Energy Transfer a donor fluorophore is excited and transfers non-radiatively energy to an acceptor molecule through dipole-dipole coupling provided that the emission curve of the donor significantly overlaps with the excitation curve of the acceptor and provided that both are less than ca. 10nm apart. The latter condition results from the dependency of the energy transfer rate (**E**) according to:

$$E = R_0^6/(r^6+R_0^6), \qquad (1)$$

where $\mathbf{R_0}$ ist the Förster distance at 50% energy transfer and **r** is the actual distance between donor and acceptor.

FRET is determined through Fluorescence Lifetime Imaging Microscopy (FLIM) and is largely independent of fluorophore concentration but sensitive to environmental factors such as temperature, viscosity and presence of fluorescence quenchers (e. g. acceptor chromophores).

The transfer efficiency (E) can also be calculated from:

$$E = 1 - \tau_{DA}/\tau_D, \qquad (2)$$

where $\boldsymbol{\tau_{DA}}$ is the fluorescence lifetime of the donor in presence of the acceptor and $\boldsymbol{\tau_D}$ is the fluorescence lifetime of the donor alone.

When FRET occurs, equation (2) gives an estimate of E, which in turn allows to calculate the molecular distance (r) by using (1) between the two interacting fluorophores fused to proteins of interest. For this $R_0$ for the two particular fluorophores is required. This was taken from the “FPbase FRET Calculator“ (https://www.fpbase.org/fret/), which estimates $R_0$ for the fluorophores EGFP and mCherry. The calculator determines $R_0$ according to:

$$R_0 = 0.211 \times \sqrt[6]{\kappa^2 n^{-4} Q_D J(\lambda)} \qquad (3)$$

where $\boldsymbol{Q_D}$ is the quantum yield of the Donor, $\mathbf{J(\boldsymbol{\lambda})}$ is the overlap integral of donor emission and acceptor excitation dependent on the wavelength $\boldsymbol{\lambda}$, $\boldsymbol{n}$ is the refractive index and $\boldsymbol{\kappa}$ is the orientation factor, taken as $\kappa^2 = 2/3$. This value for $\kappa$ is rather justified in the so called dynamic isotropic case. In this case, the angular positions of the acceptor relative to the donors emission dipole vector and on the orientation of the acceptors absorption dipole vector relative to the electric field of the donor at the acceptors location are random. Furthermore, these angular positions change rapidly during the donors excited state lifetime as a result of rapid molecular rotation. This is not given in a random isotropic population of FRET donors like GFPs fused to other proteins, which rather display a static random isotropic situation due their slow rotation. However, Vogel et al. [71] have provided a valuable approximation based on Monte Carlo Simulations for a population of potential donors and acceptors as given in a biological FRET experiment. The correction factors for low FRET efficiencies, such as the 6% in the ENP-GFP and PIN2-mCherry interaction in this study, yield almost the same approximated distance between the two fluorophores (8,25 nm) in comparison to that given in the text directly calculated from (1). The authors also indicate another approximation from R. E. Dale, which gives the same result (for discussion on the *kappa* factor see 54 and 71). The "FPbase FRET Calculator" estimates the overlap integral according to:

$$J(\lambda) = \int_0^{\infty} F_D(\lambda)\varepsilon_A(\lambda)\lambda^4\, d\lambda \,/ \int_0^{\infty} F_D(\lambda) d\lambda\ . \qquad (4)$$

where $\boldsymbol{F_D}$ is the fluorescence of the donor and $\boldsymbol{\varepsilon_A}$ is the extinction coefficient (of the solvent).

The lifetime $\boldsymbol{\tau}$ is dependent on the radiative and non-radiative rate constants according to:

$$1/\tau \ = \ k_r \ + \ k_{nr} \qquad (5)$$

According to the Strickler-Berg formula [54] the refractive index of the medium surrounding the fluorophore affects the radiative rate constant $\boldsymbol{k_r}$ and consequently the (natural) lifetime TAU/$\tau$. As seen in (3) $R_0$ also depends on the refractive index $n$. One should consider this parameter since in the “FPbase FRET Calculator“, the standard setting of $n$ is that of water (1.33). However, this parameter can diverge between different cell compartments [72, 73]. Suhling et al. [72] give a value of n = 1.35 for cytoplasm but a considerably higher value for membranes (n = 1.46 - 1.60) and have measured significantly lower lifetimes for EGFP when the refractive index n is high. For n = 1.355 (e. g. PBS with 10% Glycerol) the resulting $\tau$ is 2.68 nsec; for n = 1.463 (e. g. PBS with 90% Glycerol) the resulting $\tau$ is 2.17 nsec. The Differences for $R_0$ and E are significant between these two values. ENP is likely an associated but not an integral protein and the EGFP fluorophore is very likely exposed to the cytosol. Therefore, the refractive index value n = 1.35 (close to that of water and PBS 1.33/1.337) was taken. According to (3) the $R_0$ for the pair EGFP and mCherry is 52.36 Å (52.88 Å when n = 1.33) calculated with the “FPbase FRET Calculator“. We considered the value $R_0$ = 52.36 Å when calculating r using equation (1). Since there are no other $R_0$ data as those for EGFP and mCherry in FPbase or other reports, we used this value also for the distance calculation of GFP6 and mCherry. We did not find $R_0$ values for (E)GFP and FM4-64 and therefore cannot give an approximate distance. The occurrence of FRET between (E)GFP and FM4-64 is ascertained by comparison with the lifetime of the donor-only probe (E)GFP in the same way as for ENP-(E)GFP and PIN2-mCherry. Note, that the calculated distance is that of the fluorophores which are localized within the very center of the barrel structure of the fluorescing proteins. GFP has an (approximated) cylindrical sphere of 2nm diameter and a height of 4nm. If only considering the two fluorescent proteins, the fluorophores are 2nm apart if they are closely touching alongside and 4nm apart if they are positioned head-to-head respectively.

*Fluorescence Lifetime Imaging Microscopy*

Initially, the specimen was imaged with the parameters mentioned above using the OLYMPUS FV3000 CLSM with a 63X/NA1.2 water objective and a 488nm and a 561nm diode laser for (E)GFP and mCherry respectively. Sequential scanning of images was with a (Nyquist-adjusted) pixel size of 104nm.

FRET-FLIM of the imaged specimen was then assessed by comparing the donor lifetime (GFP construct) in absence and presence of the acceptor respectively (FM4-64, mCherry construct). We used an OLYMPUS FV3000 equipped with a Pico Quant upgrade kit including a time correlated single photon counting (TCSPC) device at a time resolution of 25psec (Time Harp 260), detection units (PMA Hybrid 40) and a laser driver (PDL 828-S SEPIAII). The GFP construct was excited with a 40MHz pulsed 485nm Laser (output 2,9µW after 60X/1.2 NA water objective, field scanning mode). The PMA Hybryd 40 units detected photons in a 520nm +/- 17.5nm window and 600nm +/- 25nm window respectively using a dicroic at 560nm. Only photon counts of the first channel (520nm +/-17.5nm) were used for GFP construct lifetime calculation with an IRF calculated by the SPT64 program (Pico Quant). The analysis results of some measurements were also analyzed after generating an IRF curve provided by Erythrosine B for comparison. Essentially, all the lifetime differences were confirmed although the Chi-square values were often inferior to those with the calculated IRF.

At pixel sizes of 104nm, photon counts of maximum 150-1000 counts (cts) for the most intensive pixels were taken. Binning was not performed. Fluorescence resolved intensity was measured for regions of interest (ROIs) embracing the apical (basal) part of epidermal (cortex) cells in the meristematic region of roots and fitted with a double-exponential re-convolution decay model using the SymphoTime64 version2.7 software

(PicoQuant SPT64; without parameter fixing). In typical scannings of such specimen only few pixels achieved the maximum count values of 150-1000 cts. However, the accumulation of all pixels representing the selected ROIs provide sufficient cts for a lifetime determination in a double-exponential case. We considered measurements, which under the applied decay model displayed a goodness of fit $Chi^2 < 1.5$, which is calculated by the SPT64 program. To obtain such $Chi^2$ values, the fluorescence decay of the EGFP and $GFP_6$ fluorophores respectively was regularly fitted with two exponentials giving two fluorescence lifetimes. We therefore always scored the average intensity lifetime $TAU_{Av\ Int}$, which is automatically calculated by SPT64 in the decay fitting process by weighting the contribution of $TAU_1$ and $TAU_2$ to the $TAU_{Av\ Int}$. The reasons for such double and multiexponential decays are complex and might be impacted by various factors such as the structure and protonation state of the fluorophore and the environment including acceptor molecules [72, 74-78]. Note that the $TAU_{Av\ Int}$. of EGFP and GFP6 are slightly different. Both proteins differ in different sequence positions. The GFP6 lacks the V in position 2, which has been introduced in many GFP versions. Both proteins possess the F64L and S65T alterations, which improve folding efficiency, fluorescence and photostability [79, 80]. The S65T alteration is also considered to suppress the (second) 395 nm excitation peak of the wild-type GFP [78, 81]. The significance of four additional differences V163A, I167T, S175G and L231H (first EGFP/second GFP6) is improved folding (V163A, I167T), neutral (L231H) or not known [81].

Analysis for significant difference were performed as Welch´s one-tailed t-Test or one-tailed t-Tests to consider unequal variances (experiments with FM4-64) or equal variances (experiments with PIN2-mCherry) respectively. All combinations showed significant differences ($p<0.0001$) except in the case of BRI1 vs. BRI1 and PIN2-mCherry.
