## Supplementary Figures for "Separate domains of the *Arabidopsis* ENHANCER OF PINOID drive its own polarization and recruit PIN1 to the plasma membrane"

### Supplementary Figures & legends

#### Overview
#### ENP structures, domains, point mutations and deletions (comparison with MEL4/NPY4)

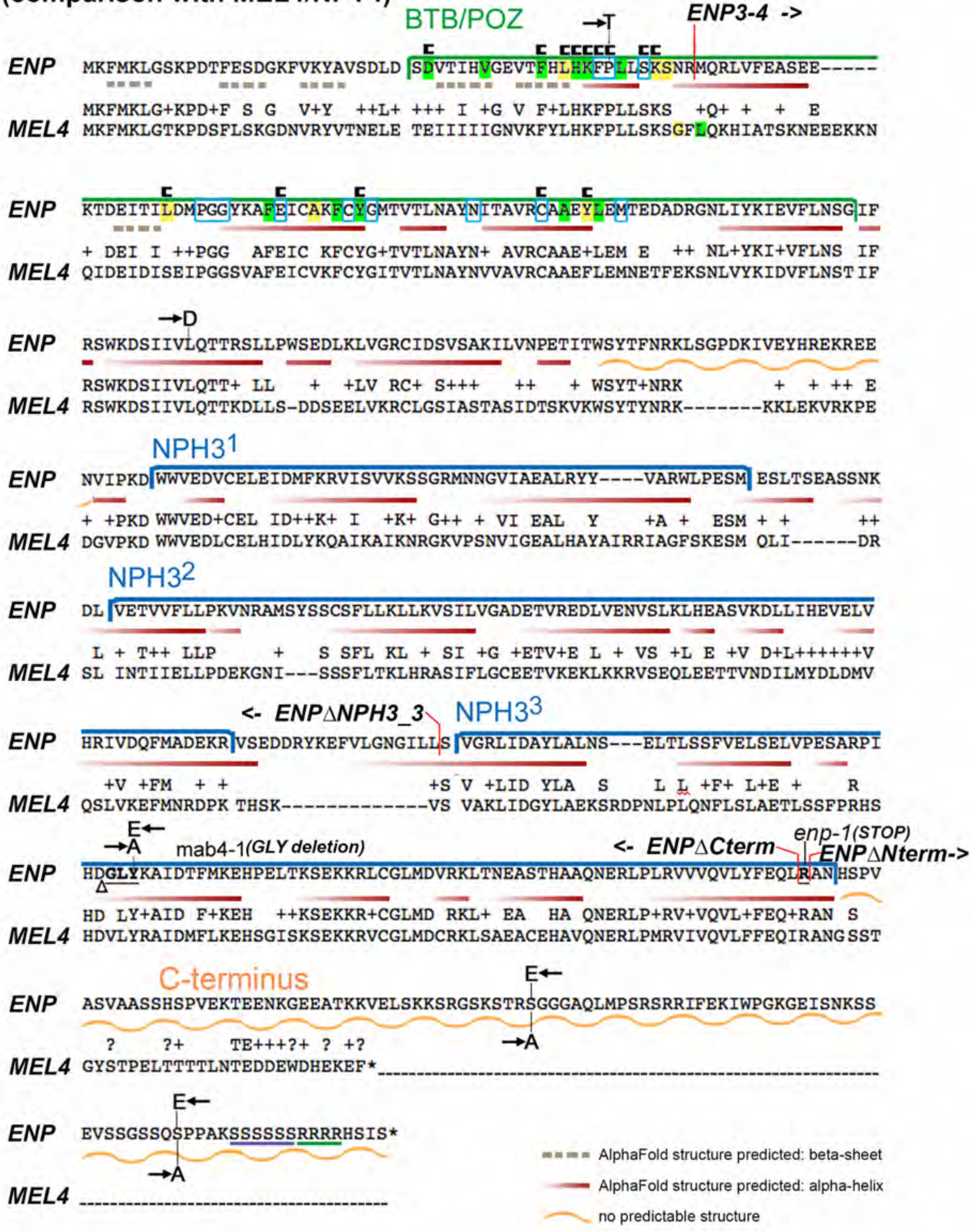


Length of ENP & MELs:

| | | | | |
|---|---|---|---|---|
| ENP | (At4g31820) | 571 aas | position | R468 100% conserved in MELs |
| MEL4 | (At2g23050) | 481 aas | homologous | R450 |
| MEL1 | (At4g37590) | 580 aas | -"- | R465 |
| MEL2 | (At5g67440) | 579 aas | -"- | R476 |
| MEL3 | (At2g14820) | 634 aas | -"- | R488 |

**SFig. 1: Overview ENP protein structure and conservation**

The Figure combines features described for the BTB/POZ domain (between aa1 and aa132) given in Stogios et al. (2005) and the structure predictions by AlphaFold (Jumper et al., 2021; Varadi et al., 2021).

The ENP sequence is compared with MEL4 (NPY4) indicating identical as well as similar ("+") aminoacids. The green bar above the aas indicates the core BTB/POZ domain (according to Stogios et al., 2005), which is then followed by sequences tentatively designated as "linker" region. The blue bars indicate the central NPH3-domain respectively, which is divided into three regions of higher similarity ($NPH3^1$ - $NPH3^3$) solely to point out the dissimilar regions in between where MEL4 is characterized by deletions.

The BTB/POZ domain includes the following information taken from Stogios et al. (2005). Yellow and green shaded aas have intermediate and high levels of conservation respectively. The latter also indicates positions that are similar in at least four of seven BTB/POZ families found eukaryotes. Aas with blue frames indicate BTB/POZ-NPH3 specific signature sequences. The c-like symbols above the (highly) conserved residues indicate those, which in four non-plant families are contact sites for known protein-protein interactions.

The different lines below the sequence indicate information given by Alphafold (Jumper et al., 2021; Varadi et al., 2021). Stippled lines in light brown indicate beta-sheet structures whereas red lines give alpha-helical structures. Shorter red lines rather represent short helical turns. Wavy yellow lines indicate a shorter (aa181-aa205) and the long C-terminal tract (aa47-aa571) of intrinsic disorder. The majority of predicted beta-sheets and alpha-helices had very high per-residue confidence metrics (pLDDT > 90). The purple and green underlining at the very end of the protein points to a potential conformation resulting from the repeated S and R motif (for details see Text).

The figure also includes all deletions of ENP analyzed in this study (the -> arrows point to the sequences retained in the truncated protein). Also added are the point mutated aas and the aas replacing them in this study. Note, that in S514 and S553 different combinations of double point mutations have been considered (see Text).

For comparison the length of ENP and all MELs is given at the bottom. The highly conserved aa R468 in the last AlphaFold predicted structure (alpha-helix) in ENP is given to indicate the length of the dissimilar C-termini of MELs beyond this site.

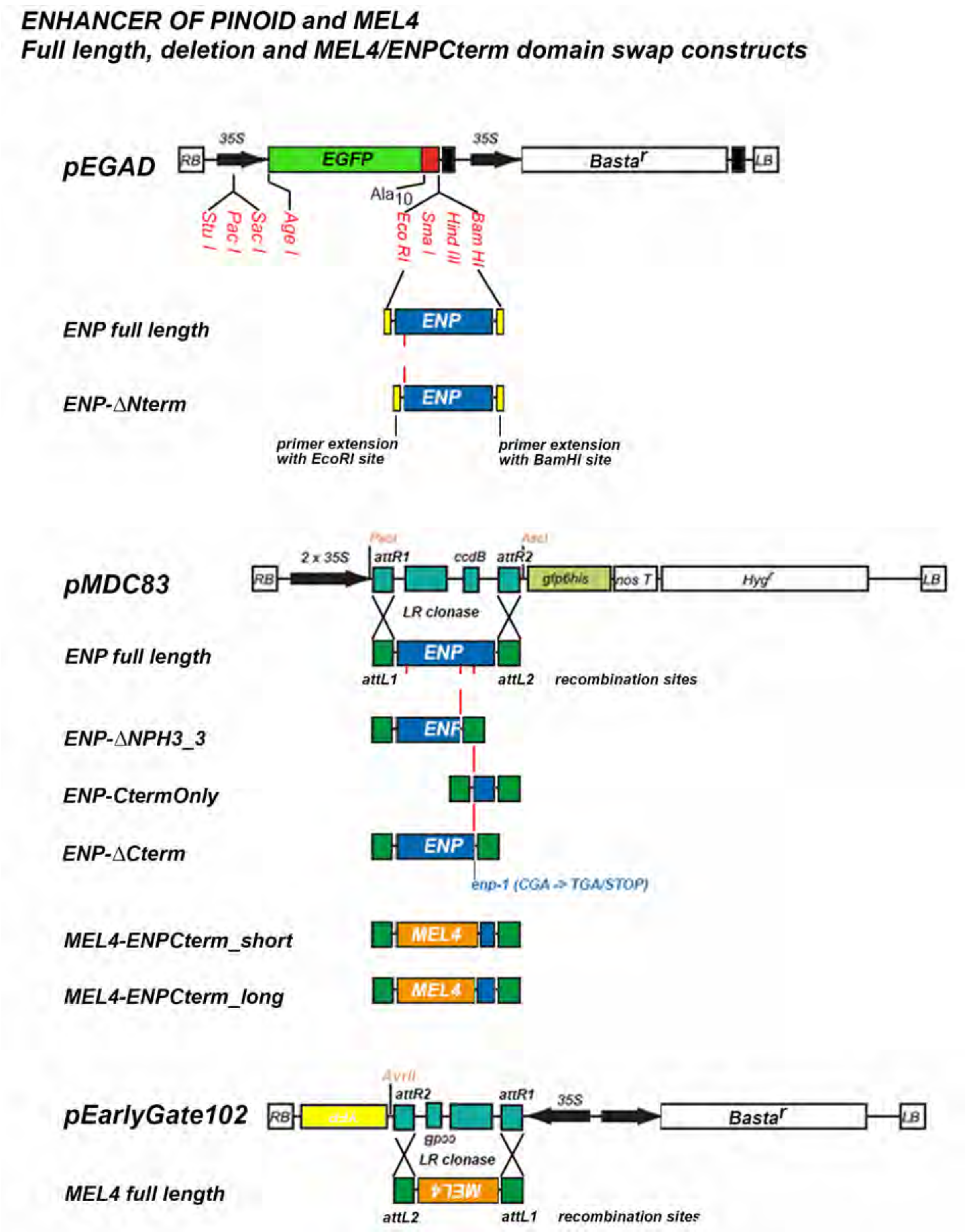


**SFig. 2: Cloning of constructs**

The figure gives a schematic (not to scale) overview of the full length and deletion constructs analyzed in this study. For N-terminal positioning via restriction-ligation of EGFP the vector pEGAD (Cutler et al., 2000) was used. All other constructs were generated via GATEWAY cloning. The ENP constructs with C-terminal GFP6 including those with point mutations were introduced into the pMDC83 vector (Curtis and Grossniklaus, 2003). The MEL4 full length construct was generated using the pEARLYGATE102 vector (for details see SText).

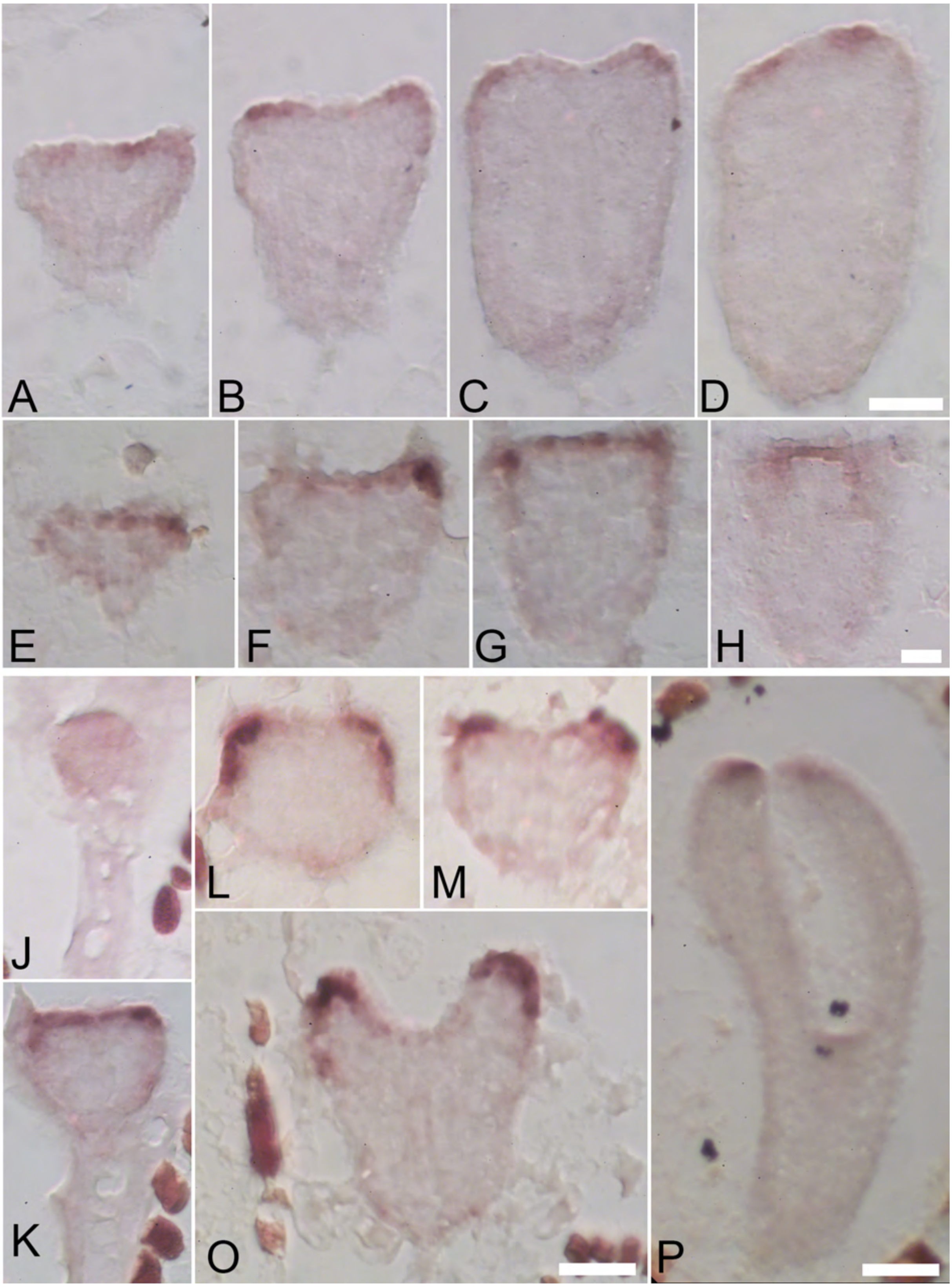


**SFig. 3: ENP expression in embryos**

In situ hybridizations with ENP antisense probe (for details see SText). A-H) Homozygous *enp pid* embryos lacking cotyledon primordia. A-D) Consecutive sections of a torpedo stage embryo with stronger ENP mRNA signal in the epidermal layer of the apex and weak signal in the root tip outer layer. E-H) Late heart stage embryo with similar expression pattern as in A-D. J-P) Wild-type embryos: 16cell-stage (J), transition stage (K), globular stage (L), early heart stage (M), late heart stage (O) and torpedo stage (P). ENP mRNA progressively concentrates from all embryo cells (J) to apical epidermis (strong) and root tip (weak)(K-O) and tips in primordia (O, P). Scale bars A-D: 20µM, E-H: 10µM, J-O: 20µM and P: 40µM.

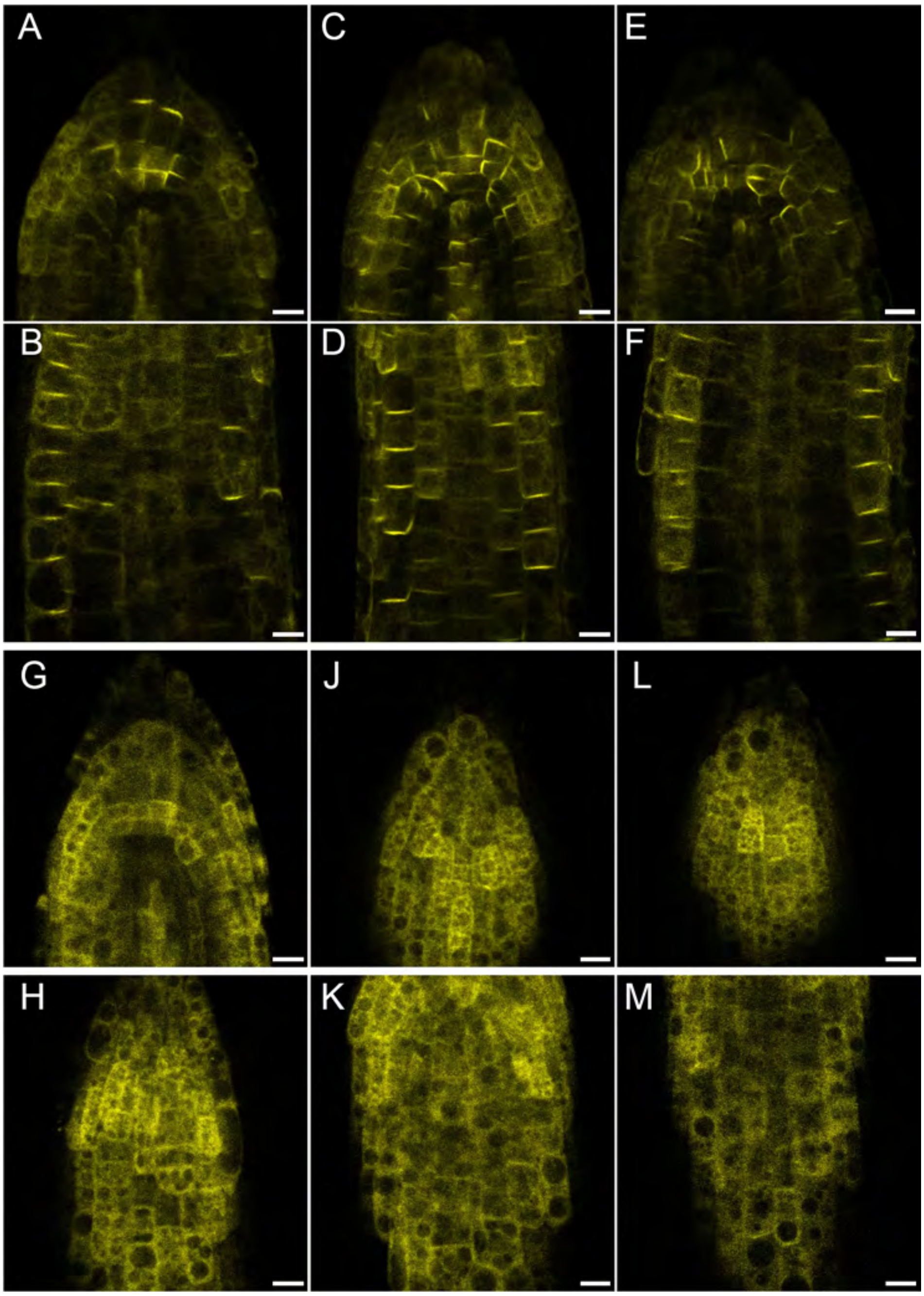


**SFig. 4: MEL4 internalization upon PBA treatment**

MEL4-EYFP shows the same response to phenylboronic acid (PBA, 10mM) as previously reported for ENP (Matthes and Torres Ruiz, 2016).

A-F) Untreated control plants: root tip (A, C, E) and the same further up (B, D, F; meristematic region, focus on epidermis). G-M) Three representative plants treated with 10mM PBA. G, J, L: root tip; H, K, M: further up. Scale bars: 10µM.

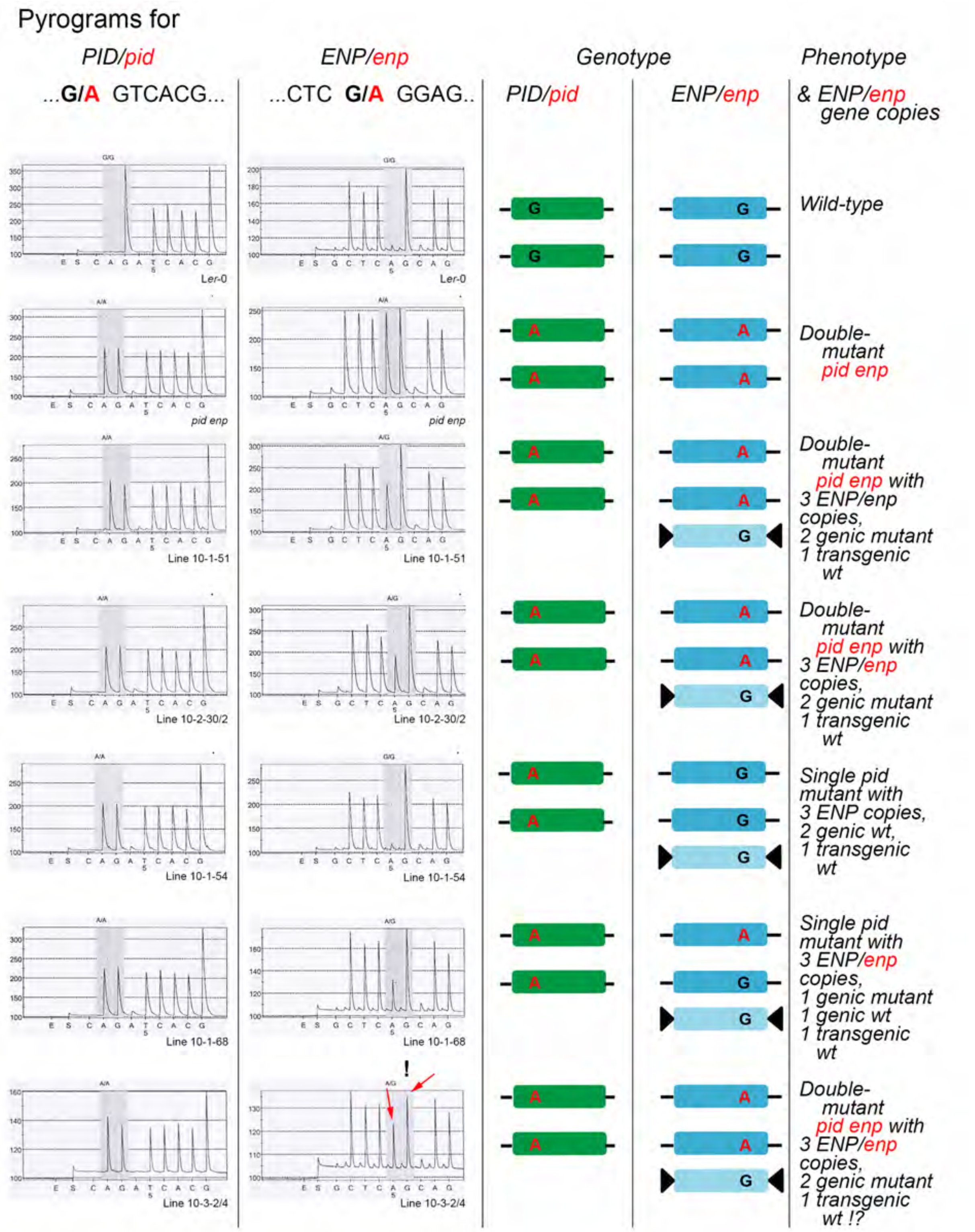


**SFig. 5: Pyrosequencing of plants.**

Critical sites, as indicated on top, of wild-type, *enp pid* double mutants and *pid* single mutants were sequenced by pyrosequencing as described (Treml et al., 2005). Point mutations sequenced were those of the *pid-15* allele (Treml et al., 2005) and *enp-1* allele (Treml et al., 2005; Furutani et al., 2007). Except the two top-most, all plants carried the full length ENP transgene *35Sp:EGFP-ENP* (symbolized by inverted black arrowheads) and were resistant to BASTA/Phosphinotricin. The phenotypes of seedlings and adult plants were either wild-type (L*er*-0; normal developed plant), *pid* (three cotyledons and abnormal but fertile flowers) and *pid enp* (no cotyledons, and rescue of flower structures). Note the height of the peaks (quantitatively) in the detection window of the pyrogram for the *PID/pid* (left) and *ENP/enp* (right) wild-type/mutant base. The bottom-most individual is a case where the height and the proportion of the peaks was not satisfying. Genotypes and phenotypes indicated.

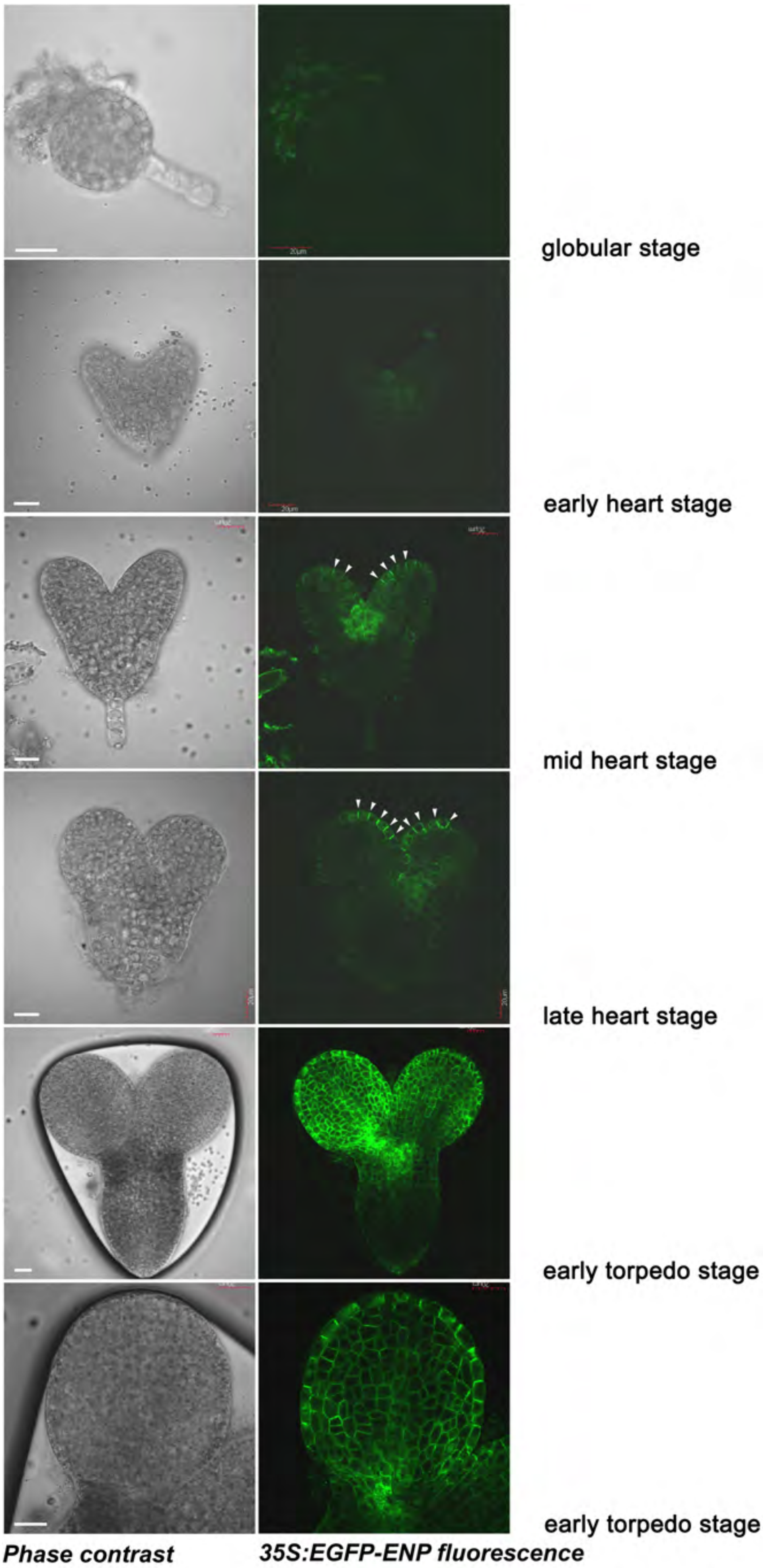


**SFig. 6: Onset of ENP expression driven by the 35S promoter**

Shown are from top to bottom as indicated: globular, early heart stage, mid heart stage, late heart stage, early torpedo stage (whole embryo and magnification of one cotyledon). A very weak signal is visible in the shoot apical meristem region in early heart stage. However as shown, cotyledon primordia do already form in this stage. In mid heart stage first polarized localization of EGFP-ENP is visible (white arrowheads). Scale bars: 20μM.

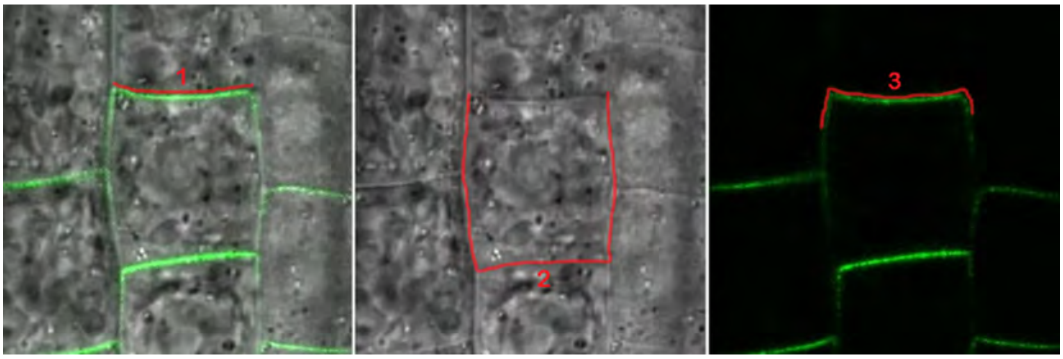


**SFig. 7: Analysis of distribution of polar ENP**

For distribution of ENP-localization within *Arabidopsis thaliana* cells, different seedlings were measured for ENP-GFP6 and ENP∆Cterm-GFP6. Only cells within the epithelial and cortex region were measured. For each cell three different measurements were made. The first for the apical membrane length (left: red line 1), the second of the residual membrane (middle: red line 2) and the third measurement for the whole length of the GFP signal (right: red line 3). The significance of differences with t-Test.

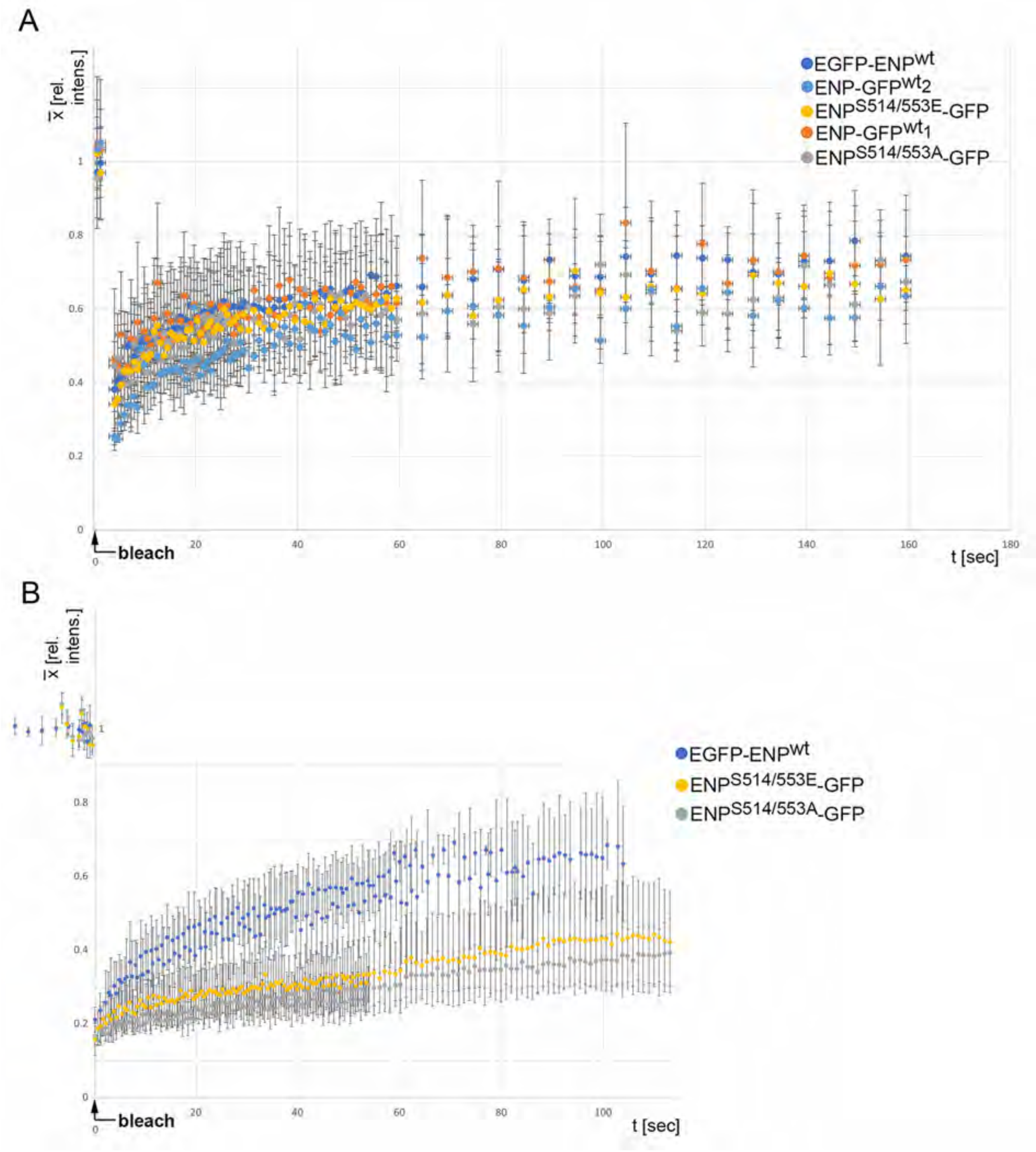


**SFig. 8: FRAP analyses of ENP constructs**

Shown is the recovery of ENP full length and double mutant ENPS514A/S553A and ENPS514E/S553E constructs applying bleaching spots of 40µm$^2$ diameter (A) and 60µm$^2$ (B). Note, that in particular the recovery dynamics of all constructs carrying the GFP6 at the C-terminus lay close together in all experiments. The recovery of the ENP full length construct carrying EGFP at the N-terminus was closer to the other constructs in the 40µm$^2$ bleach spot experiment than in the experiment with 20µm$^2$ (Fig. 3) and 60µm$^2$ diameter bleach spot. This might be due to different growth behavior of the sibling line from which the seedlings originated. Note, that the relative intensities are equal in the experiments, only the scale is shifted vertically for the intensities in B.

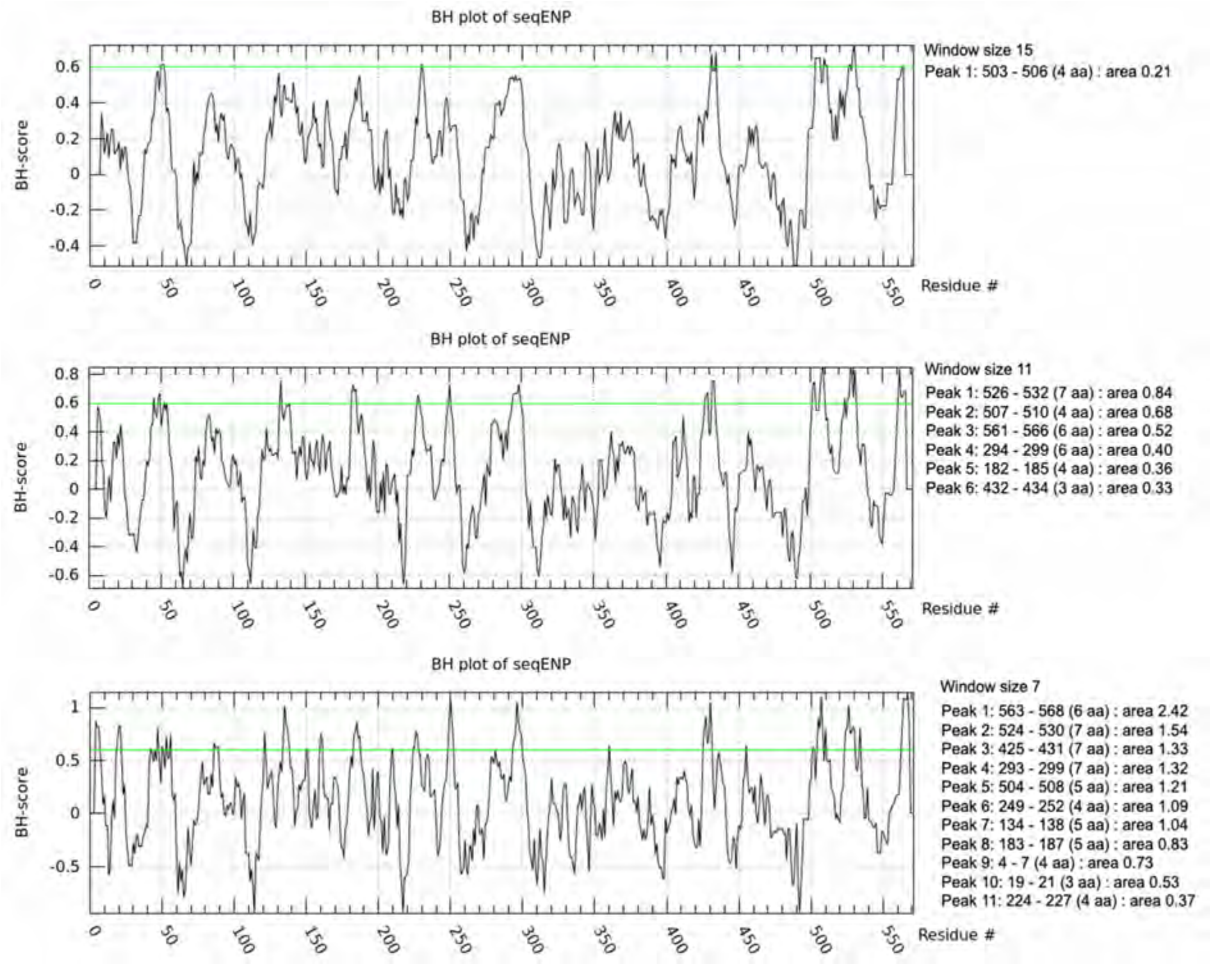


**SFig. 9: Basic and Hydrophobic plot scores of ENP**

Shown are three different Basic Hydrophobic plots (Brzeska et al., 2010) of ENP. With reduced window size, short amino acid sequence segments (from 3aas to 7 aas), along the whole ENP sequence cross the critical BH-score (green line) including the BTB/POZ, linker, NPH3-similar domain and the C-terminal IDR. Brzeska et al. (2010) explored window sizes of 19aas and 11aas to detect potential PM-associated aa segments in various genes.
